# MARS-Net: Deep learning-based segmentation pipeline for profiling cellular morphodynamics from multiple types of live cell microscopy

**DOI:** 10.1101/191858

**Authors:** Junbong Jang, Chuangqi Wang, Xitong Zhang, Hee June Choi, Xiang Pan, Bolun Lin, Yudong Yu, Carly Whittle, Madison Ryan, Yenyu Chen, Kwonmoo Lee

**Author notes:** These authors equally contributed to this work.

## Abstract

Quantitative studies of cellular morphodynamics rely on extracting leading-edge velocity time-series based on accurate cell segmentation from live cell imaging. However, live cell imaging has numerous challenging issues about accurate edge localization. Here, we develop a deep learning-based pipeline, termed MARS-Net (Multiple-microscopy- type-based Accurate and Robust Segmentation Network), that utilizes transfer learning and the datasets from multiple types of microscopy to localize cell edges with high accuracy, allowing quantitative profiling of cellular morphodynamics. For effective training with the datasets from multiple types of live cell microscopy, we integrated the pretrained VGG-19 encoder with U-Net decoder and added dropout layers. Using this structure, we were able to train one neural network model that can accurately segment various live cell movies from phase contrast, spinning disk confocal, and total internal reflection fluorescence microscopes. Intriguingly, MARS-Net produced more accurate edge localization than the neural network models trained with single microscopy type datasets, whereas the standard U-Net could not increase the overall accuracy. We expect that MARS-Net can accelerate the studies of cellular morphodynamics by providing accurate segmentation of challenging live cell images.

## Introduction

Live cell imaging is a fundamental tool to study the changes of cellular morphology (morphodynamics), which is involved in cancer metastasis, immune responses, and stem cell differentiation, among others ^1–4^. Cellular morphodynamics is governed by protrusion and retraction of the leading edges of cells, driven by cytoskeleton and adhesion processes^5, 6^. Due to their phenotypic heterogeneity, computational image analysis in conjunction with machine learning has been employed to understand and characterize cellular morphodynamics^5–8^.

Quantitative studies of cellular morphodynamics rely on extracting leading-edge velocity time-series. Therefore, accurate and consistent edge segmentation at every frame of live cell movie is necessary. Fluorescence microscopes can acquire high contrast cellular images by introducing fluorescently tagged molecules, particularly for fixed cells. Fluorescence imaging, however, causes phototoxicity to live cells, which makes researchers limit light illumination and total image acquisition time. These make fluorescence live cell images noisy, low contrast, and low-throughput. Selection of cells of low-level expression of fluorescent proteins and photobleaching further degrades the image quality^9^. Therefore, having reliable segmentation from live cell images is a significant issue. The alternative to fluorescence microscopy is a label-free phase contrast microscopy that minimizes phototoxicity in the cell. But phase contrast images contain halo and shade-off artifacts, incurring a significant challenge for reliable cell segmentation^10–13^.

Although there exist numerous conventional segmentation methods, including Otsu method^14^, Canny Detector^15^, active contour or snake-based method^16^, and PMI method based on mutual Information^17^, they often rely on a few mathematical assumptions which tend to be broken in live cell imaging conditions, rendering insufficient accuracy for cell-edge detection for the analysis of cellular morphodynamics. Supervised learning with a deep learning model can overcome such problems. Among deep learning models, Convolutional Neural Network (CNN) excels in pattern recognition in images by learning complex features directly from the input images using its hierarchical structure^18^. CNN has achieved great success in image classification^19–22^ and segmentation^23–28^ and demonstrated promising results in static and live cell images^24, 29–35^ ^36^. Particularly, U-Net^24^ is the most widely adopted CNN-based structure for image segmentation. Some of the cell image segmentation applications are easily accessible even for users without much computational resources and coding skills^35, 37, 38^. Despite these successes, the deep learning-based segmentation on live cell imaging has not been extensively tested for cellular morphodynamics studies. Moreover, models trained on their specific datasets do not generalize to different datasets, and training a new model requires large training datasets. Also, high fidelity training sets are required for training an accurate segmentation model, substantially increasing the cost of data labeling.

To increase the segmentation accuracy and overcome training data shortage for morphodynamic profiling, we developed a deep learning framework, termed MARS-Net (Multiple-microscopy-type-based Accurate and Robust Segmentation Network), which learns robust image features for accurate segmentation using the datasets from multiple types of microscopy. We reasoned that the cross-modal features learned from images of multiple types of microscopy could achieve more accurate and robust edge localization than the features from the single type of microscopy images. Therefore, we combined training data from live cell movies of migrating PtK1 cells independently taken by different microscopy techniques such as phase contrast, spinning disk confocal (SDC), and total internal reflection fluorescence (TIRF) microscopes.

In this pipeline, we used the U-Net^24^ based structure, which is CNN comprised of encoder, decoder, and skip connections in between them for segmentation. To achieve high edge localization accuracy with the datasets from multiple types of microscopy, we incorporated the transfer learning technique that initializes the weights of the network with those of the same network trained on ImageNet^39^ for the image recognition task. This has been applied in many deep learning segmentation models (FCN^23^, DeepEdge^25^, TernausNetV2^40^) and classification tasks^41–48^ to achieve high performance with a limited dataset. We replaced the U-Net encoder with one of the image classification networks such as VGG16/VGG19^20^, ResNet50V2^22^, and EfficientNetB7^49^ and used the initial weights from the ImageNet^39^ training. Among them, the pretrained VGG19 encoder coupled with U-Net decoder (VGG19-U-Net) segmented the boundary of the cell with the highest accuracy. Dropout^50^ layers were added to the model (VGG19D-U-Net) as a regularization method to prevent overfitting and boost the performance further. MARS- Net (VGG19D-U-Net trained on the images from multiple types of microscopy) was able to segment cell boundary more accurately than the model trained on single-microscopy- type data, whereas U-Net could not gain significant performance benefit from training on the data from multiple types of microscopy. Also, we demonstrate that MARS-Net enables more reliable quantitative analyses of cellular morphodynamics compared to the single- microscopy-type model.

## Results

### Overview of the computational pipeline

We prepare the ground truth masks from live cell images semi-automatically using our labeling tool (**Fig. 1a**). The images and the corresponding ground truth masks are preprocessed (see Methods for details), and they are used to train the deep neural networks for segmentation (**Fig. 1b**). The trained neural network generates a segmentation of the cell boundary, which can be used for morphodynamic profiling developed by Danuser’s group^5^ (**Fig. 1c**). It measures local velocities of the cell boundary throughout the movie and summarizes local velocities for every probing spatial window and time frame. Since this quantification method is sensitive to pixel-level segmentation errors, accurate edge localization is necessary.

**Figure 1.**
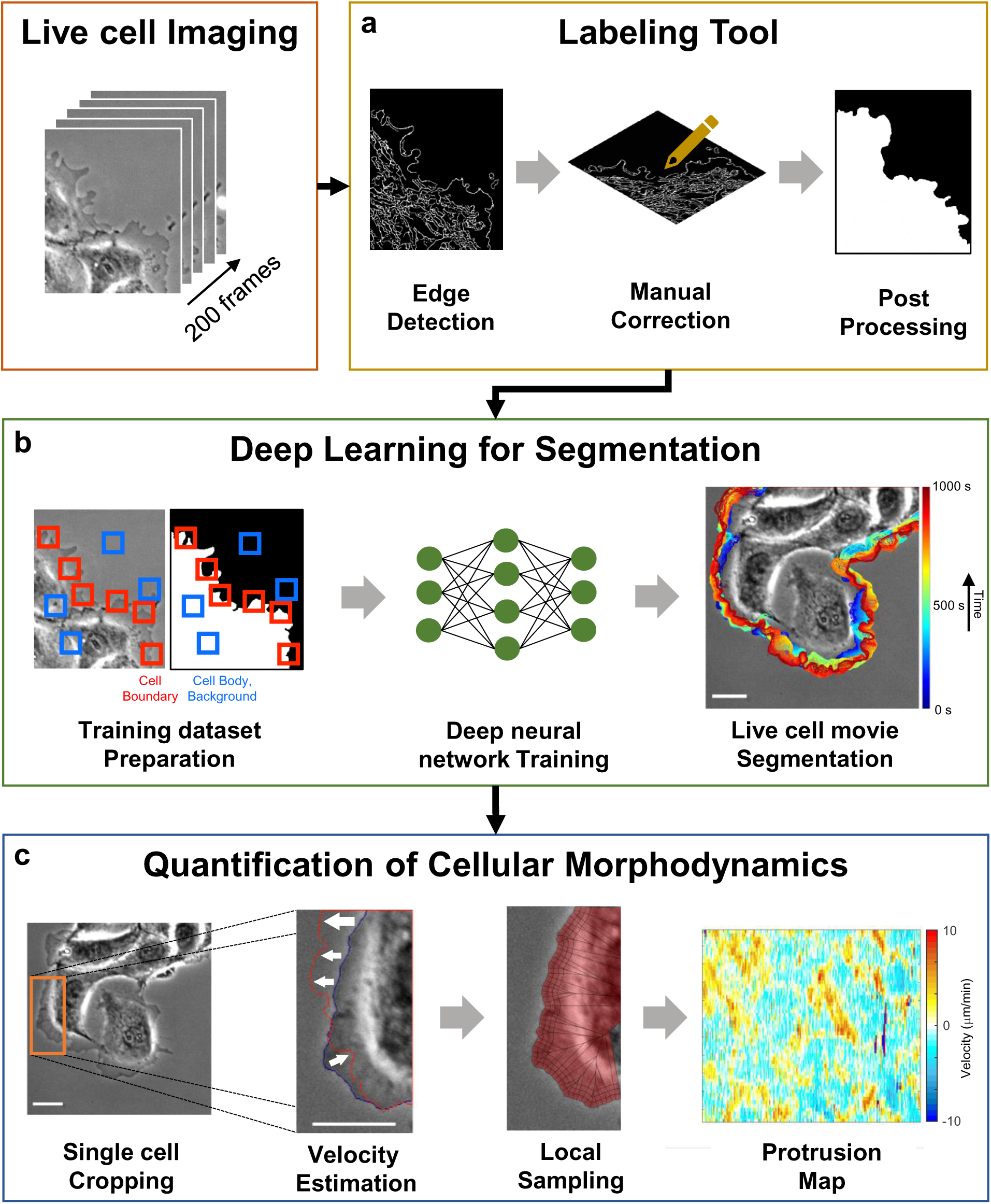
Overview of the computational pipeline. (**a**) Labeling Tool, (**b**) Deep Learning for Segmentation (MARS-Net), and (**c**) Quantification of Cellular Morphodynamics. White Bars: 32.5 μm.

Deep learning requires a large training dataset, and labeling many frames per live cell movie by hand can take several hours or days. Also, there is an inconsistency in the quality of labeled images depending on the labeler’s experience. Therefore, we created the cell labeling tool to reduce human-labor by automating most labeling procedures and reproduce accurate and consistent labels. A systematic approach to create labels promotes reliable training and evaluation of the deep learning model^29, 51^. The labeling tool takes the input image through series of image processing operations as follows. In the edge extraction step (**Fig. 2a**), the input image is blurred using Gaussian, Bilateral, and Guided blurring operations, and the Canny edge detector^15^ extracts edges from the blurred images. Three extracted edge images are combined to one edge image by adding their pixel intensity values at each coordinate. The errors such as fragmented edges and incorrect edge detection are inherent problems of conventional segmentation methods, so the users must correct the output for further processing. In the post processing step, edge images are converted into binarized segmented images, and any floating artifacts or noisy edges are removed from the edge images (**Fig. 2b**). When running the labeling tool, users have to specify which side of the edge is foreground and background and adjust two hyper-parameters based on the input image characteristics. The hyper- parameters are kernel size for blurring operations and hysteresis thresholding min-max value for detecting edges.

**Figure 2.**
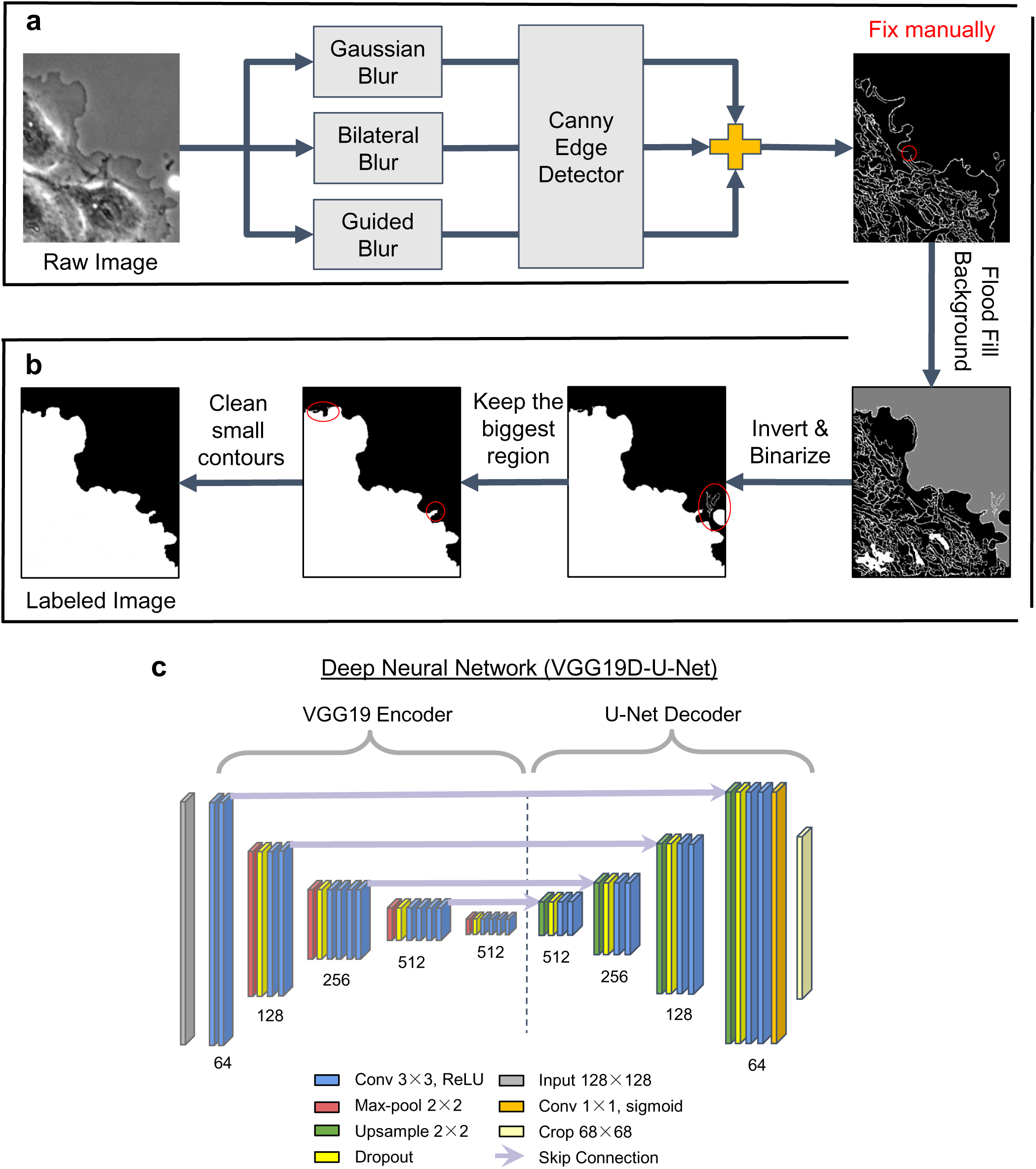
Deep learning architecture. (**a-b**) Workflow in the labeling tool comprising edge detection step (**a**) and post-processing step (**b**). (**c**) The deep neural network, VGG19D-U-Net, for segmentation of cell edges. The number of filters in each convolutional block is shown underneath each convolutional block. Light violet lines indicate which features from the encoder are concatenated with which up-sampled features in the decoder.

After the training sets are prepared, we trained the deep neural network, VGG19D-U-Net, which is a fully convolutional network with VGG19 encoder, U-Net decoder, and dropout layers (**Fig. 2c**). VGG-19 encoder contains five convolutional blocks, each of which contains one max-pooling layer and multiple convolutional layers with a depth of 64-128- 256-512-512. The first convolutional block does not have a max-pooling layer. U-Net decoder has four deconvolutional blocks comprised of one up-sampling layer that concatenates with the encoded features and two convolutional layers with the depth of 512-256-128-64. Dropout layers are added after each max-pooling and up-sampling layer. The first dropout layer is set to drop 25% of the incoming units, and the rest of the dropout layers are set to drop 50% of the incoming units.

### Segmentation of phase contrast live cell movies using VGG19D-U-Net

We first tested VGG19D-U-Net with a dataset from a single type of microscopy, which is five live cell movies of migrating PtK1 cells acquired by a phase contrast microscope for 200 frames at 5 sec/frame. The segmentation accuracy was measured by precision, recall, and F1 score of edge localization^52^ (see Methods for details) since edge localization is our main criterion for evaluation. The Wilcoxon signed-rank test was used to test the statistical significance of performance difference, unless otherwise specified. We trained models on six different numbers of training frames (1,2,6,10,22,34) from each movie. The specified number of frames were randomly selected from each live cell movie as the training data. We used the leave-one-movie-out cross validation, in which one movie is selected for testing, and the other movies are used for training.

We trained the segmentation architectures with various pretrained models integrated with U-Net decoder: VGG16, VGG19, ResNet50V2, and EfficientNetB7^53^. As demonstrated in the learning curve (**Fig. 3a**), VGG19D-U-Net converged to lower validation loss than U-Net while their training losses are the same, suggesting less overfitting in VGG19D-U-Net than U-Net. Among different encoder models, VGG19D-U-Net yielded the highest F1 score across the different numbers of training frames (**Fig. 3b**). Even with additional training frames, U-Net without pretraining could not surpass any other models trained on the equivalent number of frames. Notably, the F1 score of VGG19D-U-Net trained on one frame per movie is higher than U-Net trained on 34 frames per movie by 0.014 (0.929 vs 0.915). Overall, the F1 scores of all models tend to increase as more training frames were added, but their F1 scores plateaued as the number of training frames increased. When models were trained with ten frames per movie (**Fig. 3d-f**), F1 score of VGG19D-U-Net was significantly higher than the next best model VGG16D-U-Net by 0.003 (0.937 vs 0.934) with p=4.69x10^-6^. These results demonstrate the importance of transfer learning and dropout layers for accurate segmentation of the live cell image regardless of the size of the training dataset.

**Figure 3.**
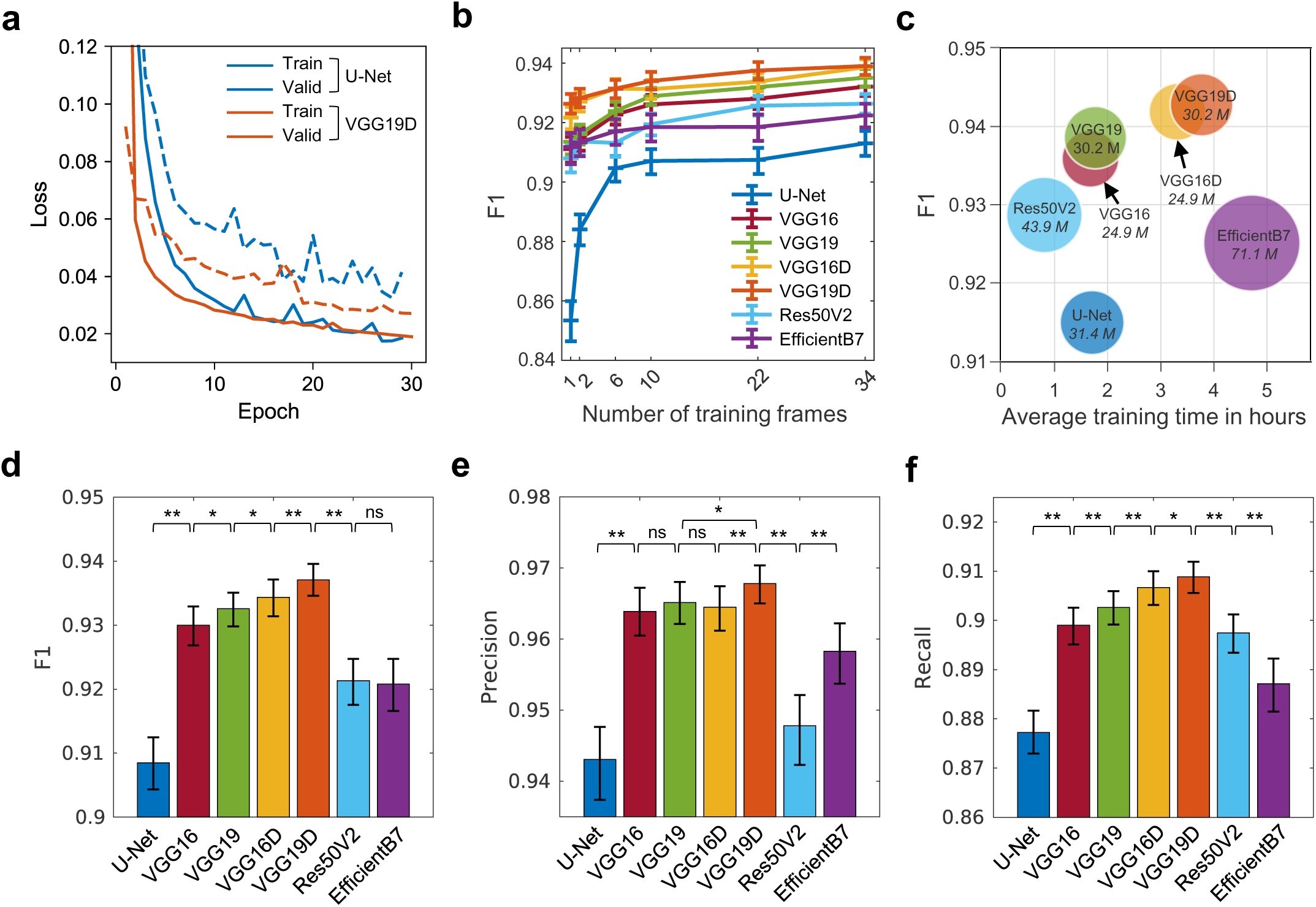
Performance comparison of the models trained on the phase contrast microscopy dataset. (**a**) Learning curves of U-Net and VGG19D-U-Net. Solid lines are average training loss, and dotted lines are average validation loss. (**b**) Average F1 scores of models trained on different numbers of frames per movie. (**c**) Training efficiency of seven different models in terms of model size, training time, and segmentation accuracy. The name of the model and its number of parameters in italics are written on the bubble. The size of a bubble is proportional to the number of parameters in the model. (**d**-**f**) Average F1, Precision and Recall of models. Suffix D denotes Dropout and suffix U-Net was omitted from the model names for brevity in the legend. The tests of significance by Wilcoxon signed-rank test with p >= 0.05 are indicated by ns, p < 0.05 are indicated by *, p < 0.0001 are indicated by **. Error bars: 95% confidence intervals of the bootstrap mean.

The model size, training time, and performance of the architectures trained on 34 frames per movie were summarized (**Fig. 3c**). The EfficientNetB7-U-Net was the deepest network with the most parameters (71.1M) and took the longest time (4.7 hours) to train on average. ResNet50V2-U-Net took the least amount of time (0.81 hours) to train but had a lower F1 than VGG16-U-Net and VGG19-U-Net. The training time among U-Net, VGG16-U-Net, and VGG19-U-Net was similar, but VGG16-U-Net and VGG19-U-net have higher F1 than U-Net. Adding dropout layers to VGG16 or VGG19 encoders (VGG16D or VGG19D) makes the model more accurate without any additional parameters but requires longer training time. Since our criterion for the best model is the high F1 score, not model size or training time, VGG19D-U-Net with the highest F1 score (0.943) was chosen as the segmentation model in our pipeline.

We also visually confirmed that VGG19D-U-Net localized the cell boundary more accurately than U-Net (**Fig. 4**). VGG19D-U-Net finds the cell body regardless of the halo effect, unlike U-Net (**Fig. 4a inset3**). Also, U-Net incorrectly segmented background as the cell body (**Fig. 4b inset1**) or segmented cell body as the background (**Fig. 4b inset2**). In the progression of the segmented cell boundary throughout the movie (**Fig. 4c**), inaccurate segmentation of U-Net in multiple frames accumulated as indicated by the white dashed box in the upper left and center of the images. In contrast, VGG19D-U-Net produced a smooth transition of the cell boundary.

**Figure 4.**
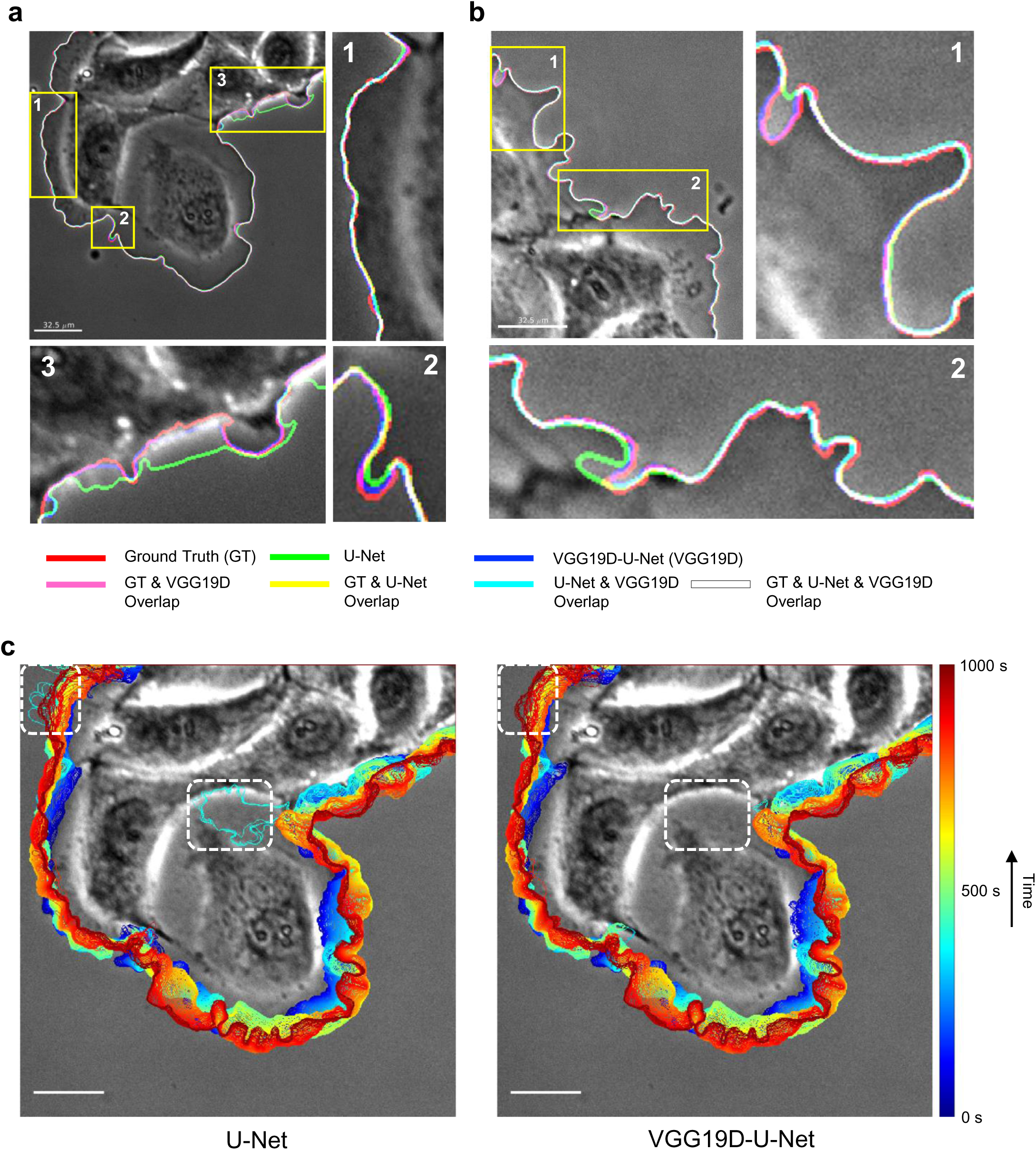
Visualization of segmentation results from U-Net and VGG19D-U-Net trained on the phase contrast microscopy dataset. (**a-b**) Edges extracted from the ground truth mask and predictions from U-Net and VGG19D-U-Net are overlaid on the first frame of the movie. Each edge is represented by one of three primary colors. Overlap of two or more edges is represented by the combination of those colors. (**c**) Progression of cell edges segmented by U-Net and VGG19D-U-Net overlaid on the first frame of the movie (blue, 0s; red, 1000s time points). Bars: 32.5 μm.

We further investigated the roles of individual components in our VGG19D-U-Net structure (**Supplementary Fig. 1a-c**). The segmentation accuracy of models in terms of F1, precision, and recall has a similar trend, so we refer to them collectively as the performance. U-Net, VGG16-U-Net, and VGG19-U-Net without pretraining have relatively similar performance. But pretraining VGG16-U-Net and VGG19-U-Net and adding dropout layers significantly increased their performance while having the same depth. Adding batch normalization layers to the VGG16-U-Net, and VGG19-U-Net reduced their performance. Also, combining a structured form of dropout for convolutional networks, DropBlock^54^, and batch normalization layers (VGG19DB-U-Net) which resembles SD- UNet^55^, resulted in significantly lower performance than VGG19D-U-Net. The performance of VGG19DB-U-Net might have been low due to the variance shift that occurs when using both dropout and batch normalization layers^56^.

Different sizes of the cropped patch are also investigated (**Supplementary Fig. 1d-f**). The size indicated here is the size of the image patches, so the size of the ground truth mask patches is smaller because our network crops out the output image (see Methods for details). The model trained on patches of size 96x96 had similar performance to other models trained on bigger patches. The models trained on 128x128, 192x192, and 256x256 patches had almost identical precision (0.969). However, if the size of a patch was reduced to 80x80 or 64x64, the performance of the model decreased significantly. These results suggest that the features relevant to detecting cellular boundary exist in the patch size of 128x128 even though it lacks contextual information of the entire cell body. Since training on smaller patches has the benefits of reducing memory usage and training on more diverse patches using the same computational resources, we used the patch size of 128x128 in our pipeline.

### Segmentation of live cell movies from a single type of fluorescence microscopy using VGG19D-U-Net

In this section, we tested the segmentation accuracy of VGG19D-U-Net using fluorescence live cell movies. The training sets consisted of five live cell movies of PtK1 cells expressing GFP-mDia1 using a spinning disk confocal (SDC) microscope for 200 frames at 5 sec/frame, and six live cell movies of PtK1 cells expressing paxillin-HaloTag-TMR acquired by a Total Internal Reflection Fluorescence (TIRF) microscope for 200 frames at 5 sec/frame. These live cell images are very challenging for the segmentation using conventional intensity-thresholding methods. The SDC images are highly noisy and low contrast because the cells expressed low levels of GFP-mDia1. Although the TIRF images have higher contrast and less noise than SDC images, they have other technical challenges as follows: i) high-intensity signals of paxillin accumulated in focal adhesions make edge segmentation difficult, particularly for intensity threshold-based methods, ii) the nonuniform light illumination of a TIRF microscope incurs additional issues for the segmentation, iii) the leading edge of cells could transiently lift up and leave the thin TIRF illumination, resulting in less visible cell edges.

To prepare the reliable segmentation training sets for the SDC images, we also expressed SNAP-tag-actin and label it by TMR (SNAP-tag-TMR-actin) and performed the multiplexed imaging together with GFP-mDia1. The images in the channel of SNAP-tag- TMR-actin have good contrast along the cell boundary. Therefore, conventional image thresholding was applied to SNAP-tag-TMR-actin images, and the resulting binary masks were used as ground truth labels for the SDC datasets. To make more reliable ground truth masks for the TIRF images, the fluorescence images of the same cells were also taken using standard widefield illumination. We used our labeling tool to label the cell edges in the widefield images, which served as ground truth labels for the TIRF datasets.

We trained U-Net and VGG19D-U-Net on the fluorescence SDC and TIRF datasets separately, and evaluated their performance by leave-one-movie-out cross validation. During training (**Fig. 5a,d**), VGG19D-U-Net converged to lower validation loss than U-Net while U-Net overfitted as epoch increased, as demonstrated by the increase of difference between training and validation loss. Across the different number of training frames, VGG19D-U-Net yielded a higher F1 score than U-Net on SDC datasets (**Fig. 5b**). For one of the movies, U-Net performed considerably worse than VGG19D-U-Net, indicated by the faint line with the least F1 score of about 0.2. Even though the average F1 score seems to stay consistent as the number of training frames increased in the graph, VGG19D-U-Net trained on 34 frames per movie had a greater average F1 score than the same model trained on one frame per movie by 0.028 (0.891 vs 0.863). When models were trained on ten training frames per movie, VGG19D-U-Net had a greater F1 than U-Net by 0.115 (0.866 vs 0.751) with p=2.95x10^-16^. Also, the distribution of evaluated frames in the F1 score (**Fig. 5c**) showed that all frames which were evaluated as 0 in F1 score when segmented by U-Net, had higher F1 scores when segmented by VGG19D. The superior performance of VGG19D-U-Net compared to U-Net is consistent with the results on phase contrast datasets.

**Figure 5.**
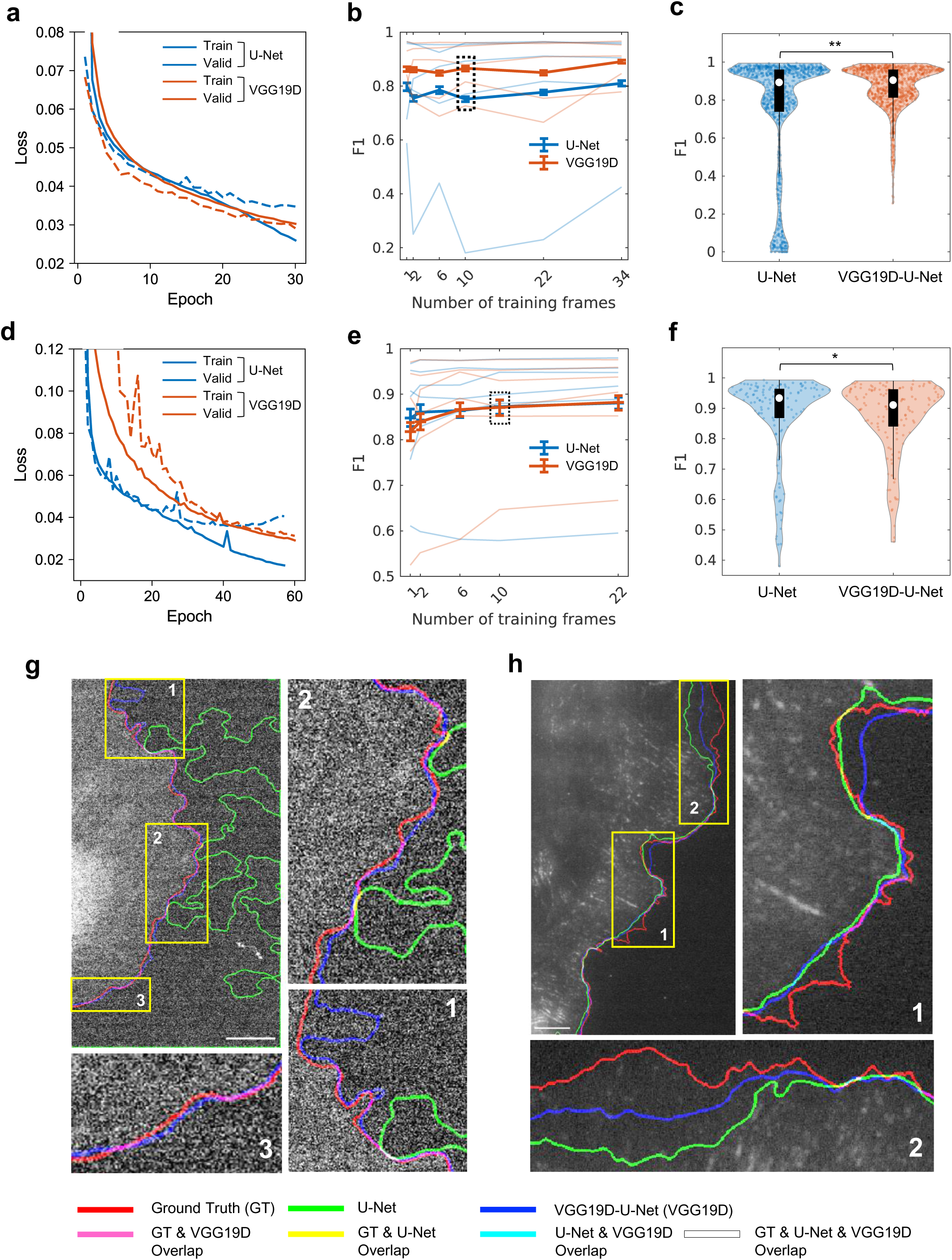
Performance comparison of U-Net and VGG19D-U-Net trained on fluorescence microscopy datasets. (**a-c, g**) Models trained on SDC datasets and (**d-f, h**) Models trained on TIRF datasets. (**a,d**) Learning curves of U-Net and VGG19-U-Net. Solid lines are average training loss, and dotted lines are average validation loss. (**b, e**) Average F1 scores of models trained on varying numbers of training frames. Lighter lines represent individual test set results in the leave-one-movie-out cross validation, and darker and thicker lines represent the average of all test set results. Error bars: 95% confidence intervals of the bootstrap mean. (**c, f**) The distribution of F1 score in violin plot and box plot in black with a median indicated by the white circle. Number of evaluated frames are (**c**) n=1000 and (**d**) n=132. The tests of significance by Wilcoxon signed-rank test with p < 0.05 are indicated by * and p < 0.0001 are indicated by **. **(g-h)** Visualization of edges extracted from ground truth masks and predictions from U-Net and VGG19D-U- Net overlaid on the first frame of the movie. Each edge is represented by one of three primary colors. Overlap of two or more edges is represented by the combination of those colors. (**g**) Bar: 7.2 μm. (**h**) Bar: 6.5 μm.

On TIRF datasets (**Fig. 5e**), U-Net initially surpassed VGG19D-U-Net when trained on 1 or 2 frames per movie, but they converged to similar average F1 scores as the number of training frames increased. When models were trained on two frames per movie, VGG19D-U-Net had a lower average F1 score than U-Net by 0.02 (0.840 vs 0.860) with p=2.88x10^-7^. But when models were trained on ten training frames per movie, VGG19D- U-Net had a marginally lower average F1 score than U-Net by only 0.002 (0.871-0.873) with p=0.002. This similarity is also reflected in the distributions of the evaluated frames of both models (**Fig. 5f**).

The visual inspection of edges segmented by U-Net and VGG19D-U-Net demonstrated that VGG19D-U-Net performed well on all SDC datasets, while U-Net failed to segment one of the SDC datasets (**Fig. 5g**). Both U-Net and VGG19D-U-Net did not perform well on one of the TIRF datasets, shown by the mismatch between the ground truth edge and segmented edges from U-Net and VGG19D-U-Net (**Fig. 5h**). The mismatch is because of the limited illumination of the cell boundary that is hard to detect even for human eyes. The ground truth mask was created using the same cell images by widefield illumination, so a portion of the cell edge may be lifted from the surface and away from the thin illumination of the TIRF microscope.

### Training of VGG19D-U-Net on the datasets from multiple types of microscopy

We established that VGG19D-U-Net outperformed U-Net when they were trained with the phase contrast and the SDC datasets, respectively. Now, we prepare the training set from multiple types of microscopy by combining the previous data (phase contrast, SDC, and TIRF microscopy) and train VGG19D-U-Net on them to create MARS-Net. Same as the evaluation for the single-microscopy-type models, leave-one-movie-out cross validation was used. When this training strategy was applied, multiple-microscopy-type VGG19D- U-Net (VGG19D-U-Net^M^, MARS-Net) converged to lower validation loss than multiple- microscopy-type U-Net (U-Net^M^) without overfitting (**Fig. 6a**). VGG19D-U-Net^M^ had a significantly higher F1 than single-microscopy-type VGG19D-U-Net (VGG19D-U-Net^S^) by 0.028 (0.904 vs 0.876), whereas there was not a significant difference in F1 between U- Net^M^ and single-microscopy-type U-Net (U-Net^S^) (**Fig. 6b**). The distribution of the evaluated frames (**Fig. 6c**) from VGG19D-U-Net^M^ also contained fewer outliers than other models.

**Figure 6.**
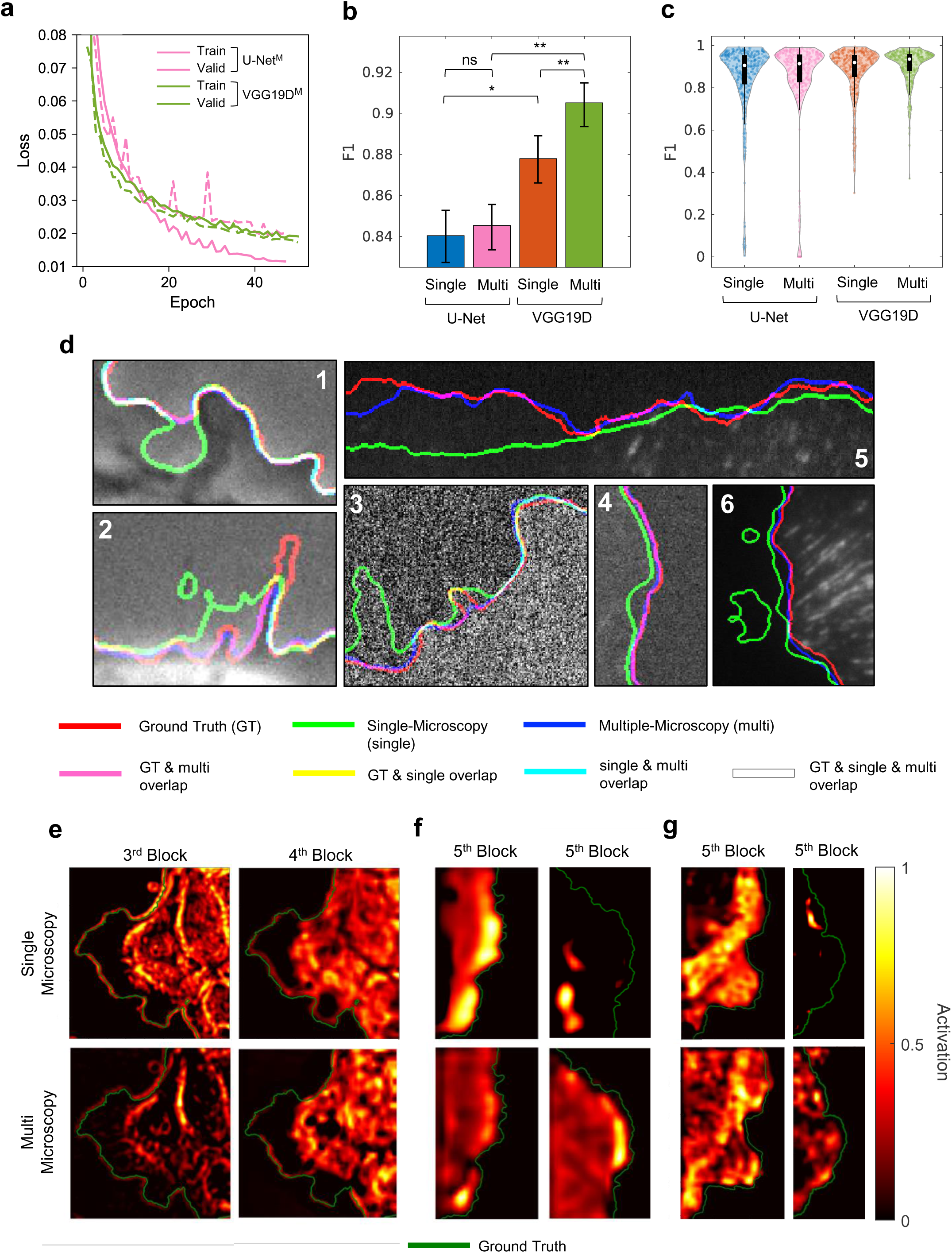
Comparison of single-microscopy-type and multiple-microscopy-type training using U-Net and VGG19D-U-Net. (**a**) Learning curves of U-Net^M^ and VGG19- U-Net^M^. Solid lines are average training loss, and dotted lines are average validation loss. (**b**) Average F1 scores across all datasets. “Single” represents single-microscopy- type, “Multi” represents multiple-microscopy-type, and “VGG19D” represents the VGG19D-U-Net. The statistical significance by Wilcoxon signed-rank test with p >= 0.05 are indicated by ns, p <0.05 are indicated by *, and p < 0.001 are indicated by **. Error bars: 95% confidence intervals of the bootstrap mean. (**c**) The distribution of F1 scores in the violin plot and the box plot in black with a median indicated by the white circle. The statistical significances are not shown since they are indicated in (**b**). The number of evaluated frames is n=335. Every fifth frame in the movie is sampled to gather about 21 frames from each movie. (**d**) Close-up views of the segmentation results on all three microscopies: phase contrast, SDC and TIRF. Edges extracted from ground truth masks and predictions are overlaid on their corresponding original image. Each edge is represented by one of three primary colors. Overlap of two or more edges is represented by the combination of those colors. (**e-g**) Class activation map of the single-microscopy- type and multiple-microscopy-type VGG19D-U-Net with respect to the ground truth edge. The last layer in the third and fourth block of the encoder is visualized for one randomly chosen frame in the phase contrast (**e**) live cell movie in order from left to right. The last layer in the fifth block of the encoder is visualized for one randomly chosen frame in two of the SDC (**f**) and TIRF (**g**) live cell movies. In the heatmap, the value 0 means no activation, and 1 means the highest activation. The green line represents the ground truth edge of the cell body.

When the performance was averaged per microscopy type (**Supplementary Fig. 2 a-c**), VGG19D-U-Net^M^ had a higher F1 than VGG19D-U-Net^S^ for every microscopy type; It had marginally significantly higher F1 in the phase contrast dataset by 0.003 (0.933 vs 0.930) with p=0.039 (the paired sample t-test was used since their differences in F1 are normally distributed according to the Lilliefors test (p=0.062)). Also, VGG19D-U-Net^M^ significantly improved F1 than VGG19D-U-Net^S^ in the SDC dataset by 0.05 (0.911 vs 0.861) with p=2.61x10^-51^ and in the TIRF dataset by 0.029 (0.878 vs 0.849) with p=0.012 by Wilcoxon signed-rank test. While U-Net^M^ marginally improved F1 than U-Net^S^ in the SDC dataset by 0.012 (0.767 vs 0.755) with p=1.39x10^-16^ and not significantly improved F1 in the TIRF datasets with p=0.068, U-Net^M^ significantly reduced F1 for the phase contrast dataset by (0.884 vs 0.898) with p=1.43x10^-5^. The distributions of the evaluated frames (**Supplementary Fig. 2d-f**, **Supplementary Fig. 3**) show that MARS-Net can accurately segment many frames that VGG19D-U-Net^S^ could not handle in the SDC and TIRF datasets.

In addition, the performance of MARS-Net was similar or greater than U-Net^M^ on all microscopy types. Unlike VGG19D-U-Net^S^, which had a significantly lower F1 than U- Net^S^ in the TIRF dataset, the difference of F1 between MARS-Net and U-Net^M^ was not significant (p=0.637). The distributions of the evaluated frames (**Supplementary Fig. 2d-f**, **Supplementary Fig. 3**) show that MARS-Net can accurately segment outlier frames that U-Net^M^ could not handle in the phase contrast and SDC datasets. These results demonstrate that our deep learning architecture, VGG19D-U-Net was more effective in learning the cross-modal features from the datasets of multiple types of microscopy and generalizing to unseen datasets than U-Net.

We also visually confirmed the performance between VGG19D-U-Net^S^ and MARS-Net in edge localization of all three microscopies (**Fig. 6d**, **Supplementary Fig. 2g-i**). In most cases, both accurately localizes the edge and overlaps with the ground truth illustrated by white lines. However, in the cases that VGG19D-Net^S^ made inaccurate edge localization, MARS-Net accurately localized the ground truth edge, shown as pink lines. Even for one of the TIRF movies that VGG19D-U-Net^S^ struggled in the previous section (**Fig. 5h inset2**), MARS-Net can localize edges more accurately (**Fig. 6d inset5**). Taken together, VGG19D-U-Net can be trained with live cell images from multiple types of microscopy and produce more accurate and robust segmentation than the single- microscopy-type model.

To understand the effect of multiple-microscopy-type training on VGG19D-U-Net, we made the class activation maps of the convolutional layers in the encoder using SEG- GRAD-CAM^57^ (**Fig. 6e-g**). The class activation map shows which pixels in the original image positively influence the feature maps in the convolutional layer to segment the cell boundary pixels. In the encoder of VGG19D-U-Net comprised of five blocks of convolutional layers, the last layers in each block are visualized. In phase contrast images, the activation map from the third and fourth block showed consistent differences in activated features between single and multiple-microscopy-type models (**Fig. 6e**). In the third block, the multiple-microscopy-type model utilized features both on the outside and inside of the edge, while the multiple-microscopy-type model mainly utilized features on the outside of the edge exclusively for segmenting cell boundary. The activated regions from the multiple-microscopy-type model are associated with the brightness outside the cell boundary due to the halo effect in phase contrast microscopy. Also, in the fourth block, the multiple-microscopy-type model mainly utilized features inside the cell boundary, while the multiple-microscopy-type model utilized features along the cell boundary. These results illustrate that how changing the training dataset influence the same deep learning model to utilize different features in the image for segmentation.

Complete arrangement of class activation maps from first to last blocks in the encoder for all live cell movies (**Supplementary Fig. 5-7**) demonstrates that the earlier block spots low-level, fine-grained features, and the last block spots the entire cell body. Based on this observation, the multiple-microscopy-type models could spot the whole cell body in fluorescence images (SDC and TIRF), while the multiple-microscopy-type model could only find a portion of the cell body for some of the movies (**Fig. 6f-g**). In total, the multiple- microscopy-type model could not find the entire cell body for 4 out of 11 fluorescence movies, while the multiple-microscopy-type model correctly found cell bodies in all fluorescence movies. These results suggest that the cross-modal features learned from the dataset from multiple types of microscopy are more effective than the multiple- microscopy-type dataset.

**Figure 7.**
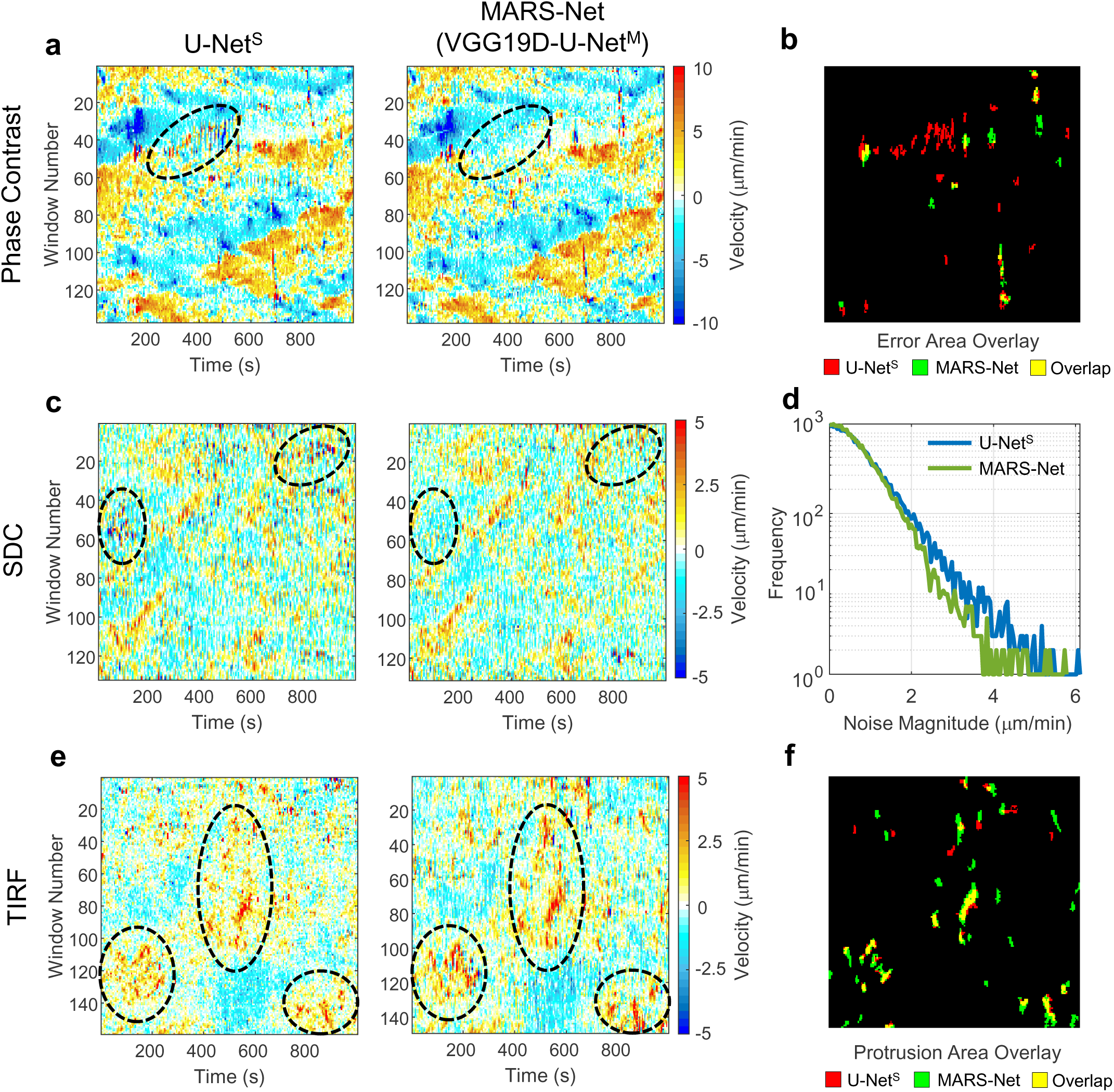
Morphodynamic profiling of the segmented movies by single- microscopy-type U-Net and MARS-Net. (**a**, **c**, **e**) Protrusion velocity maps made from U-Net^S^ and MARS-Net. The black ellipses emphasize some of the erroneous regions (**a,c**) from U-Net^S^ and the improved protrusion patterns from MARS-Net (**e**). (**b**) Overlay of the large regions of error/noise containing at least 10 pixels in the protrusion maps of the phase contrast movie (**a**) from U-Net^S^ and MARS-Net. (**d**) Comparison of overall noise present in protrusion maps of the SDC movie (**c**) from U-Net^S^ and MARS-Net. (**f**) Overlay of the large protrusion areas containing at least 10 pixels in the protrusion map of the TIRF movie (**e**) from U-Net^S^ and MARS-Net.

### Quantitative Profiling of Cellular Morphodynamics

After segmenting phase contrast, SDC and TIRF live cell movies by U-Net^S^ and MARS- Net, we quantified local protrusion velocities to see how MARS-net improves cellular morphodynamics profiling^5^ over the standard U-Net. The protrusion maps of phase contrast movies segmented by MARS-Net contain less errors or noise than the protrusion maps by U-Net^S^ (**Fig. 7a**). We define the velocity errors as the dramatic change in protrusion or retraction velocity within a few frames due to the segmentation error, indicated by alternating red and blue color. To corroborate these visual observations, we located the errorneous parts in the protrusion map by thresholding the noise images from each protrusion map (**Fig. 7b**, **Supplementary Fig. 8**) (see Methods for details). For the phase contrast movie, when pixels in large errorneous regions are counted, U-Net^S^ produced more errors than MARS-Net by (567 vs 307). In the SDC movie, U-Net^S^ not only produced more errorneous regions than MARS-Net but also produced more background noise (**Fig. 7c**). When all magnitude of noise were counted and plotted against their frequency, U-Net^S^ produced more noise than MARS-Net (**Fig. 7d**). For the TIRF movie, instead of reducing the noise, MARS-Net exhibited stronger patterns of protruding cell boundary than U-Net^S^, illustrated by long columns of red in the black ellipses (**Fig. 7e**). For the quantitative comparison, we thresholded the noise-filtered protrusion maps (**Fig. 7f**) . When the pixels in large regions of the protrusive regions were counted, MARS-Net produced more protrusion than U-Net^S^ by (1013 vs 582). Strongly protruding edges have low contrast since they are lifted upward and away from the TIRF illumination. Our analysis demonstrates that MARS-Net is capable of detecting these edges, allowing more accurate morphodynamic profiling. Taken together, considering protrusion maps can be used to identify phenotypes from subcellular movement^7^, both error/noise reduction and protrusion enhancement of the morphodynamics pattern from the accurate segmentation of MARS-Net will benefit further analysis.

## Discussion

Our transfer learning approach employing VGG19 and dropout layers is shown to be superior to conventional U-Net for segmenting live cell time-lapse images in both single- microscopy-type and multiple-microscopy-type training. ImageNet pretrained VGG19-U-Net has been effective in the segmentation of medical images^58, 59^. Also, VGG19 encoder were better than other encoders with deeper layers, Res50V2 and EfficientNetB7. This may be because ResNetV2 and EfficientNetB7 reduce the input spatial dimension by half in their first convolutional layers, while VGG19 encoder preserves the input spatial dimension with convolutional layers that does convolution with padding. Since our objective is to segment cell boundary accurately, retaining low-level features in the first convolutional layers that can identify edges^60^ are crucial for the localization of cell boundary. Moreover, the labeled training images required to get satisfactory segmentation results on the dataset from multiple microscopy types are only 1% (two frames per movie of 200 frames) in the leave-one-movie-out cross validation, which can facilitate cell biologists to adopt MARS-Net in their research.

Cell images from different microscopies can have drastically different image qualities and distribution of intensities, so training on them together could have degraded the performance by confusing the model instead. In the similar work done by CellPose^35^, multiple types of cell images were combined to build one generalist model. Its generalist model had a similar segmentation accuracy to the specialist model trained on one type of cell image when they were evaluated on the specialist dataset. On the contrary, training on live cell movies of three different microscopes (phase contrast, SDC, TIRF), we were able to significantly enhance the segmentation model by extracting effective features for cell edges across different microscopy data instead of overfitting on single microscopy data. Remarkably, although three types of live cell images employed in this study are very difficult to be segmented by conventional algorithms, the cross-modal features synergistically learned by MARS-Net were able to successfully detect extremely low contrast cell edges that could not be detected by the single-microscopy-type model due to the noise and the limited TIRF illumination (**Figure 6d, inset5 and 6**)

Through transfer learning with ImageNet pretrained weights, the deep learning model reuses diverse features learned from millions of images on the Internet^39^. This benefits the model to become invariant to various imaging conditions such as brightness, contrast, and camera resolution. Similarly, multiple-microscopy-type training benefits VGG19D-U-Net to become invariant to the imaging modality and attempt to create a robust model that identifies cell boundaries with semantic understanding, as shown by our activation maps. Another benefit of multiple-microscopy-type training is that it reduces the need to create new training datasets because the training dataset in one microscopy can be reused to analyze the dataset in another microscopy. This is consistent with the previous study demonstrating that multi-fidelity data was used to increase the size of the training set and improved the performance of the deep learning model in material science research^61^.

Morphodynamic profiling has been usually done with high-contrast fluorescence live cell images amenable for standard threshold-based segmentation methods, limiting the throughput of the analysis pipeline. Particularly, phase contrast microscopy images had not been used due to the segmentation issues. Since phase contrast microscopy does not require expensive optical components and fluorescence labeling, MARS-Net, in conjunction with phase contrast microscopy, can substantially accelerate quantitative studies of cellular morphodynamics.

## Methods

### Data Collection

Cell culture and live cell imaging procedures were followed according to the previous studies^7^. PtK1 cells were cultured in Ham’s F12 medium (Invitrogen) supplemented with 10% FBS, 0.1 mg ml^−1^ streptomycin, and 100 U ml^−1^ penicillin. Cells were then imaged at 5 second intervals for 1000 seconds using 0.45 NA Super Plan Fluor ELWD 20X ADM objective for phase contrast imaging and 60X, 1.4 NA Plan Apochromat objective for fluorescence spinning disk confocal imaging, 1.49NA Apochromat TIRF 100X for fluorescence TIRF imaging.

PtK1 cells were transfected with the DNA constructs of GFP-mDia1 and SNAP-tag-actin or paxilin-HaloTag using Neon transfection system (Invitrogen) according to the manufacturer’s instructions (1 pulse, 1400 V, 20 ms) and were grown on acid-washed glass #1.5 coverslips for 2 days before imaging. Prior to imaging, expressed SNAP-tag-actin or paxillin-HaloTag proteins were labeled with SNAP-tag-TMR (New England BioLabs) or HaloTag-TMR (Promega) ligands, respectively according to the manufacturers’ instructions. All imaging was performed in imaging medium (Leibovitz’s L- 15 without phenol red, Invitrogen) supplemented with 10% fetal bovine serum (FBS), 0.1 mg ml^−1^ streptomycin, 100 U ml^−1^ penicillin, 0.45% glucose, 1.0 U ml^−1^ Oxyrase (Oxyrase Inc.) and 10 mM Lactate. PtK1 cells were acquired from Gaudenz Danuser lab. They were routinely tested for mycoplasma contamination.

All microscopy except for TIRF microscopy (described elsewhere^6^) was performed using the set up as follows: Nikon Ti-E inverted motorized microscope (including motorized focus, objective nosepiece, fluorescence filter turret, and condenser turret) with integrated Perfect Focus System, Yokogawa CSU-X1 spinning disk confocal head with a manual emission filter wheel with Spectral Applied Research Borealis modification, Spectral Applied Research custom laser merge module (LMM-7) with AOTF and solid state 445 nm (200 mW), 488 nm (200 mW), 514 nm (150 mW), 561 nm (200 mW), and 637 nm (140 mW) lasers, Semrock 405/488/561/647 and 442/514/647 dichroic mirrors, Ludl encoded XY stage, Ludl piezo Z sample holder for high speed optical sectioning, Prior fast transmitted and epi-fluorescence light path shutters, Hamamatsu Flash 4.0 LT sCMOS camera, 37 °C microscope incubator enclosure with 5% CO2 delivery (In Vivo), Molecular Devices MetaMorph v7.7, TMC vibration-isolation table.

### Dataset

The live cell movies used for training and evaluation of our pipeline are as follows.

• Five movies of label-free migrating PtK1 cells by a phase contrast microscope
• Five dual-color movies of PtK1 cells expressing GFP-mDia1 and SNAP-tag-actin by a Spinning Disk Confocal (SDC) microscope.
• Six movies of PtK1 cells expressing paxillin-HaloTag-TMR, a marker of cell-matrix adhesions by a Total Internal Reflection Fluorescence (TIRF) microscope

Each live cell movie contains 200 frames, and about 40 frames per movie were labeled by our labeling tool for each phase contrast movie. For each SDC movie, all 200 frames were labeled by thresholding actin images. For each TIRF movie, 22 frames were labeled using the images from standard widefield fluorescence microscopy images and our labeling tool. Overall, 202 frames from phase contrast, 1000 frames from SDC, and 132 frames from TIRF movies are labeled to train and test our pipeline. The pixel size is 325nm for phase contrast datasets, 72nm for SDC datasets and 65nm for TIRF datasets.

The ground truth masks for phase contrast and TIRF images are labeled using our labeling tool (**Fig. 2a-b**). SDC images have the corresponding high contrast images of SNAP-tag-TMR-actin with good contrast along the cell boundary. Therefore, ground truth masks for SDC images can be labeled by applying denoising and thresholding without human intervention. The non-local means method implemented in ImageJ for denoising (sigma=15 and smoothing_factor=1) was applied to each SNAP-tag-TMR-actin image. Then, thresholding was applied to all frames with an optimal threshold determined by visually checking the generated masks and re-adjusting the threshold until the generated masks align the best with the cell boundary. The generated binary masks were used as the ground-truth for GFP-mDia1 fluorescence videos.

### Training Dataset Preparation

Before training the deep learning model, frames and their ground truth mask are processed for training and testing. Six different numbers of training frames (1,2,6,10,22,34) are randomly selected from each live cell movie except for the test set movie to determine the adequate number of training frames to train the model. The chosen frames become part of the training/validation set. Then, 200 patches of 128X128 pixels are randomly cropped from each frame and its corresponding ground truth mask. The cropping is necessary to reduce memory and computational requirement and to ensure that the size of the input image works in our model. 60% of the cropped images are from the boundary of cytoplasm illustrated by red boxes, and 20% are from inside, and other 20% are from outside of the cytoplasm illustrated by blue boxes in **Fig. 1b**. Both cropped images and masks are in grayscale and called patches. Only patch sizes in multiple of 16 are allowed because other sizes cause a mismatch of spatial size between encoded features and decoded features when concatenating them in U-Net structure.

Patches are augmented to negate the effect of small training size and improve performance. The augmentation methods include random rotation within 50 degrees, width, and height shift within 10% of the image’s width and height, shear in counter- clockwise direction within 36 degrees, zoom in or out randomly within 10% of the image size, and horizontal and vertical flips of the image. The original image’s reflection replaces the portion of the augmented image outside the boundary of the original image. The default number of augmented patches is 6400. For instance, the total number of patches in training and validation sets given two frames from each live cell movie in leave-one- movie-out cross validation is 8000 (2 x 4 x 200 + 6400). Then, patches are randomly split into training and validation sets with a ratio of 80:20.

Image patches are preprocessed to facilitate the deep learning model training. For phase contrast and SDC datasets, all image patches from one movie are standardized based on the mean µ and the standard deviation δ of pixel values of the cropped and augmented patches in that movie. In this way, the distribution of pixel values per movie has the mean and standard deviation equal to zero and one, respectively. Image patches from the TIRF dataset have poor contrast, so they are preprocessed differently from phase contrast or SDC datasets. After the mean µ and the standard deviation δ of pixel values of a TIRF movie are calculated, the pixel values *x*_*i,j*_ are replaced with the following values when they are less than µ − 2δ or greater than µ + 3δ.

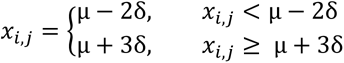

In the end, the min-max normalization was applied to rescale the pixel ranges to [0, 1].

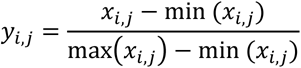

For prediction, images and masks are not cropped or augmented, but the same standardization or preprocessing steps are applied based on the microscopy type.

### Neural Network Architecture

All models mentioned in this paper are based on the same U-Net structure comprised of encoder and decoder. The original U-Net encoder was replaced by other encoders such as VGG16, VGG19, ResNet50V2, and EfficientNetB7, but the same U-Net decoder was used without any modification. Weights of the standard U-Net encoder were randomly initialized, and encoders in other models were pretrained with ImageNet provided by Keras. Every model had four skip connections that concatenate encoded features with decoded features.

The convolutional filter size is 3x3, and the zero-padding in each convolutional layer of U-Net and VGG models yields the feature map with the same spatial size after convolution. The size of the input patch is 128x128, and the size of the max-pooling and up-sampling filter is both 2x2. The same size of max-pooling and up-sampling filters make the max-pooled feature map and the up-sampled feature map at the same hierarchical level to have the same spatial size. The last layer of the network crops the 128x128 output by 30 pixels on all sides to get the segmented image of size 68x68. Cropping is necessary to eliminate the boundary effects. Without cropping, the segmented image is hazy along the image boundary, lowering the segmentation model’s accuracy.

In the prediction step, every frame of the movie in the test set is segmented by the trained model. The image is not cropped into 128x128 pixel patches, but its width and height are padded with its reflection by about 30 pixels. Then, the padded regions are removed from the predicted binary mask to avoid boundary effects.

### Neural Network Training Settings

For a fair comparison of models, each model’s hyperparameters were configured the same as follows: Adam^62^ optimizer with learning rate=10^-5^, batch size=64, input size=128, output size=68, early stopping patience=3. In the decoder and encoder that is not pretrained, the kernel weights were initialized with Glorot uniform^63^, and the bias weights were initialized with zeros. The pretrained models were fine-tuned without freezing any weights.

The binary cross-entropy was used as a loss function for training. To avoid overfitting, we used the early stopping, so training stopped when the validation loss did not decrease during the three consecutive epochs. For the phase contrast and SDC datasets, early stopping patience was 3, and the maximum epoch was 100. For the TIRF dataset, early stopping patience was 10, and the maximum epoch was 300. We used default parameters in the Keras for other parameters. The neural network training was performed using TensorFlow^64^ 2.3 on RTX Titan GPU with CUDA 10.1 for multiple-microscopy-type phase contrast models and multiple-microscopy-type models and TensorFlow 1.15 on GTX 1080Ti GPU with CUDA 10.0 for multiple-microscopy-type SDC models and multiple-microscopy-type TIRF models.

### The Number of Training Frames

The number of training frames per movie in leave-one-movie-out cross validation (F_C_) and the total number of training frames (F_T_) used to train the model in the results section are described here.

• F_C_ = 34, F_T_ = 136 (**Fig. 3c**)
• F_C_ = 10, F_T_ = 40 (**Fig. 3a,d-f , Fig. 5a,c,g, Supplementary Fig. 1**)
• F_C_ = 10, F_T_ = 50 (**Fig. 5d,f,h**)
• F_C_ = 2, F_T_ = 8 (**Fig. 4**)
• F_C_ = 2, F_T_ = 30 (**Fig. 6a**)
• F_C_ = 2, and F_T_ = 30 for multiple-microscopy-type models, F_T_ = 8 for single- microscopy-type model trained on phase contrast or SDC datasets or F_T_ = 10 for single-microscopy-type model trained on TIRF dataset (**Fig. 6b-g, Fig. 7 Supplementary Fig. 2-7**).

### Cross Validations

To rigorously test our deep learning model’s generalizability and reproducibility, we evaluated every model by leave-one-movie-out cross validation. It is the same as leave-one-subject-out cross validation^65^ but with the subject replaced by the live cell movie. We set aside one movie as a test set, and the rest of the movies are used for training and validation. Frames in the same live cell movie have little difference in image features, but there is a distinctive visual difference even among the live cell movies taken by the same microscopy. Therefore, we consider frames in the same live cell movie to be independent and identically distributed (i.i.d) and live cell movies to be out of distribution (o.o.d). The leave-one-movie-out cross validation ensures that our model is assessed on the o.o.d test set to prevent shortcut learning^66^. For instance, given five live cell movies called A, B, C, D, and E, movie E is set aside as a test set, and movies A, B, C, and D become training/validation sets. In the subsequent validation, movie D is set aside as a test set, and movies A, B, C, and E become training/validation sets. This process is repeated until every movie is set aside as the test set once. Then, the test performance measures are averaged.

Our segmentation pipeline trained on the same dataset can yield different segmentation results due to random selection of frames, random cropping, and random train/validation set split. In order to reduce the variations caused by them, the leave-one-movie-out cross validation is repeated five times for single-microscopy-type phase contrast models in **Fig. 3**, and **Supplementary Fig. 1a-c**. Frames and patches were randomly selected and randomly split into training and validation set in each repetition.

### Evaluation Metrics

Precision, recall, and F1 score between ground truth edges and segmented edges are calculated by the edge correspondence algorithm in the Berkeley Segmentation Benchmark^52, 67^, with the search radii (Phase Contrast: 3 pixels; SDC: 5 pixels, TIRF: 5 pixels). The package performs bipartite matching of two edge images by iteratively matching how edge pixels in one image corresponds to edge pixels in another image. For instance, an edge pixel in the first image is counted as a match if there is an edge pixel in the second image within the search radii of the target pixel.

Before evaluation, both ground truth masks and segmentation by the models are image processed. The image processing steps include thresholding the grayscale images into binary images with a threshold value of 0.5, given intensity values ranging from 0 to 1, filling small holes, and extracting edge by the canny edge detector. Since images are binarized before evaluation, the intensity value of each pixel in the image is either 0 or 1. The match between ground truth pixels and segmented pixels of intensity 1 are true positives (*tp*). The segmented pixels of intensity 1 that do not match with ground truth pixels are false positives (*fp*). And the ground truth pixels of intensity 1 that do not match with segmented pixels are false negatives (*fn*). True negatives, which are the match between ground truth pixel of intensity 0 and segmented pixel of intensity 0, are ignored. After counting *tp*, *fp*, *and fn* in an image, the metrics are calculated as follows.

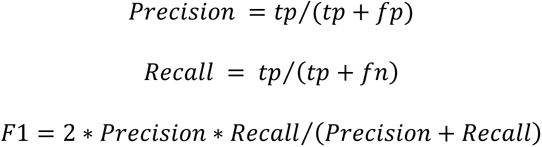

For each metric, every segmented frame is evaluated, and evaluated frames are bootstrapped with 1000 replicates to calculate bootstrap mean and 95% confidence interval. The image processing and evaluation are done using MATLAB R2019b.

### Profiling of Cellular Morphodynamics

The steps taken to perform quantitative profiling of cellular morphodynamics^5^ on segmented movies is described in detail (**Fig. 1c**). The live cell movie is cropped as illustrated by the dashed white rectangle, and the velocity of the cell along its boundary is estimated based on the difference of segmented area in the previous and present frames. Then, the estimated velocity is grouped into rectangular blocks called "window" to get a smoother estimate of the velocity. At the local sampling step, the outermost band of windows along the boundary of the cell is sampled to draw a protrusion activity map showing the velocity at each window number and frame number. Inner bands inside the cell are ignored. The size of each window is 6 pixels, or 1.95μm for phase contrast dataset, 7 pixels, or 504nm for SDC, and 8 pixels, or 520nm for TIRF datasets.

The signal in the protrusion map is found by the cubic smoothing spline interpolation of the protrusion maps with the smoothing parameter (p=0.7). The error/noise is calculated by subtracting the original protrusion map with the spline filtered protrusion map. To ignore the regions of error/noise and protrusion of small magnitudes, thresholding operation set magnitude lower than 3μm/min to zero for the phase contrast movie and lower than 2μm/min to zero for the SDC and the TIRF movies. Then, small connected regions of error/noise or protrusion signal containing less than 10 pixels are removed (**Fig. 7b,f** and **Supplementary Fig. 8**) to highlight the large error from U-Net^S^ or protrusion signal from MARS-Net that facilitates further analysis. For the SDC dataset (**Fig. 7d**), the histogram of all error magnitude without thresholding was plotted against its log frequency as a line graph.

### Class Activation Map

The technique called SEG-GRAD-CAM^57^ visualizes the feature maps that are positively associated with the increase in the intensity of output pixels. Unlike GRAD-CAM^68^, which is designed for classifiers that output a vector, SEG-GRAD-CAM can explain the decision of the segmentation model that outputs a 2-dimensional matrix. Our region-of-interest is the cell boundary, so we visualized activation of feature maps with respect to the edge extracted from the ground truth mask.

Let A^k^ be the k^th^ feature map in the filter. Among convolutional layers in the VGG19D-U-Net, we are interested in the last layer of each block in the encoder. The total number of feature maps from the first to last blocks are as follows: 64, 128, 256, and 512. The output of the model, *y,* only has one channel, and its value ranges from 0 to 1. *i* and *j* are indexes of the pixels that correspond to the cell boundary *C* in the output image *y*, and *u* and *v* are indexes of the spatial location in *A^k^*. N is the total number of pixels in *A^k^*. Then, the importance of the feature map at each spatial location can be computed as follows.

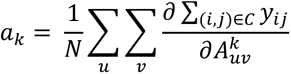

For every pixel of *y*, the gradients with respect to all pixels in the feature map are calculated by backpropagation and global average pooled across the spatial dimensions of *A^k^*. The weight matrix *ak*, which the same spatial dimension as *A^k^*, dot products with *A^k^* to get the weighted sum of the feature maps.

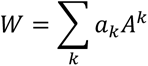

The weighted sum of the feature maps or heatmap is spatially scaled up by bilinear interpolation to match the input image size. Scaling up is necessary because the heatmaps from different convolutional feature maps have different spatial sizes. Scaled-up heatmaps are overlaid with their corresponding ground truth edge and can be compared with each other. Finally, ReLU is applied to ignore the negative influence of the feature maps on the prediction of a cell boundary.

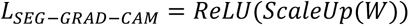

## Code availability statement

The code for our deep learning-based segmentation pipeline and evaluation code is available at https://rc-gitlab.chboston.org/kleelab/MARS-Net

## Data availability statement

The datasets used in the current study are available from the corresponding author on a reasonable request.

## Acknowledgments

We thank Microsoft for providing us with Azure cloud computing resources (Microsoft Azure Research Award), and Boston Scientific for providing us with the gift for deep learning research. This work was supported by NIH grant GM122012 and GM133725.

## Author Contributions

CW and XZ initiated the project. JJ and CW designed the pipeline. JJ wrote the final version of the manuscript and supplement. JJ, CW, XZ, BL, and YY wrote the code for the segmentation model. JJ built the labeling tool. HC performed the live cell imaging experiments. JJ, CW, HC, XP, MR, and YC prepared the training sets. JJ visualized feature maps using SEG-GRAD-CAM and profiled cellular morphodynamics from the segmented movies. KL coordinated the study and wrote the final version of the manuscript and supplement. All authors discussed the results of the study.

## Author Information

Correspondence and requests for materials, data, and code should be addressed to KL.

---

**Supplementary Figure 1.**
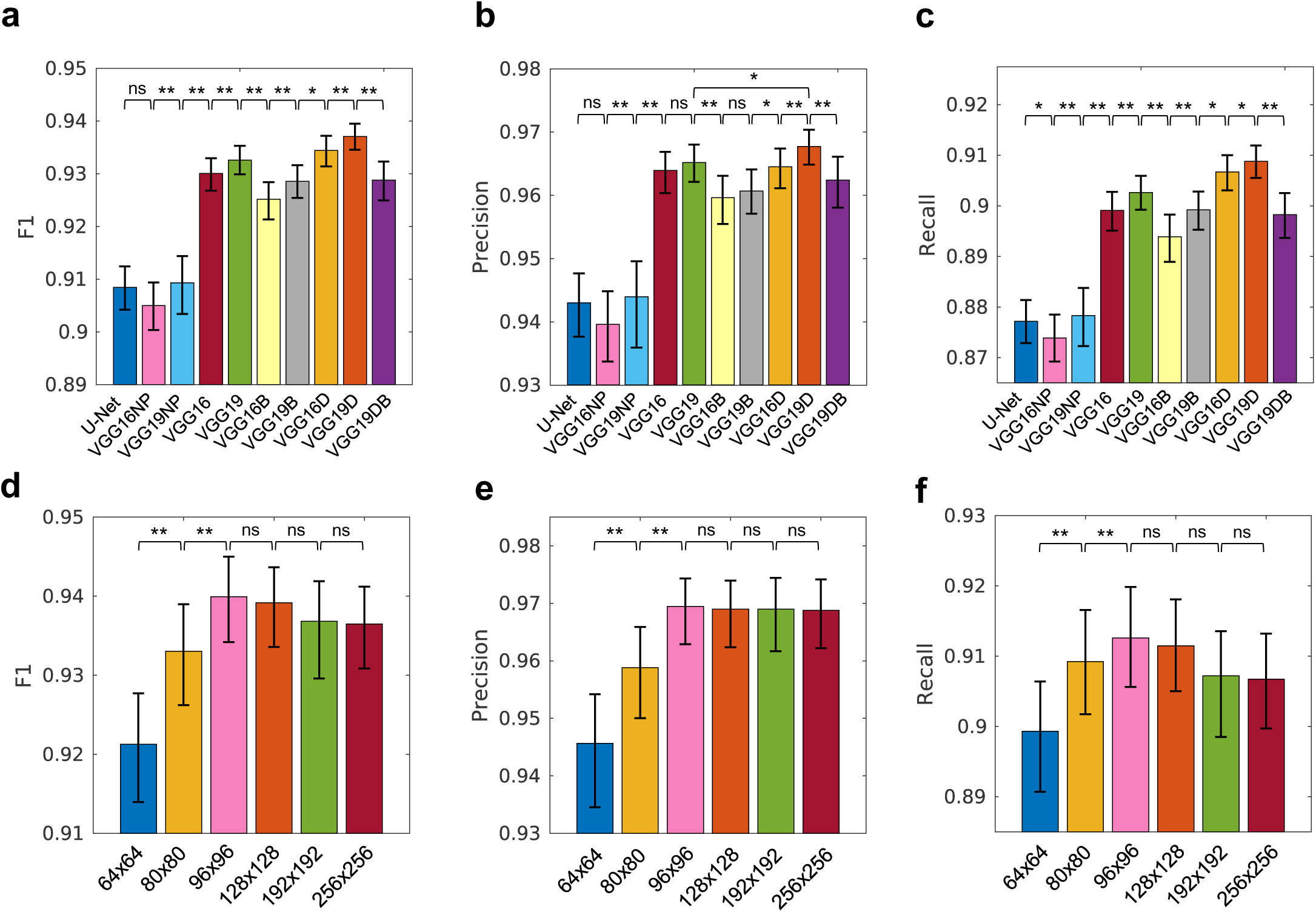
Effects of VGG variants and patch size on the models trained on phase contrast datasets. (**a-c**) Average performance of U-Net and its variants with VGG encoders, pretraining, batch normalization or dropout layers. Model names suffixed by “NP” and U-Net are not pretrained, but other models are ImageNet pretrained. Suffix B denotes batch normalization and D denotes Dropout. (**d-f**) Average performance of ImageNet pretrained VGG19D-U-Net trained on different sizes of two-dimensional cropped image patches, ranging from 64x64 to 256x256 pixels. The tests of significance by Wilcoxon signed-rank test with p >= 0.05 are indicated by ns, p < 0.05 are indicated by * and p < 0.0001 are indicated by **. Only the statistical tests of differences between adjacent bars are shown except for (**b**) to compare VGG19D-U-Net with the next best model, VGG19-U-Net. Error bars: 95% confidence intervals of the bootstrap mean.

**Supplementary Figure 2.**
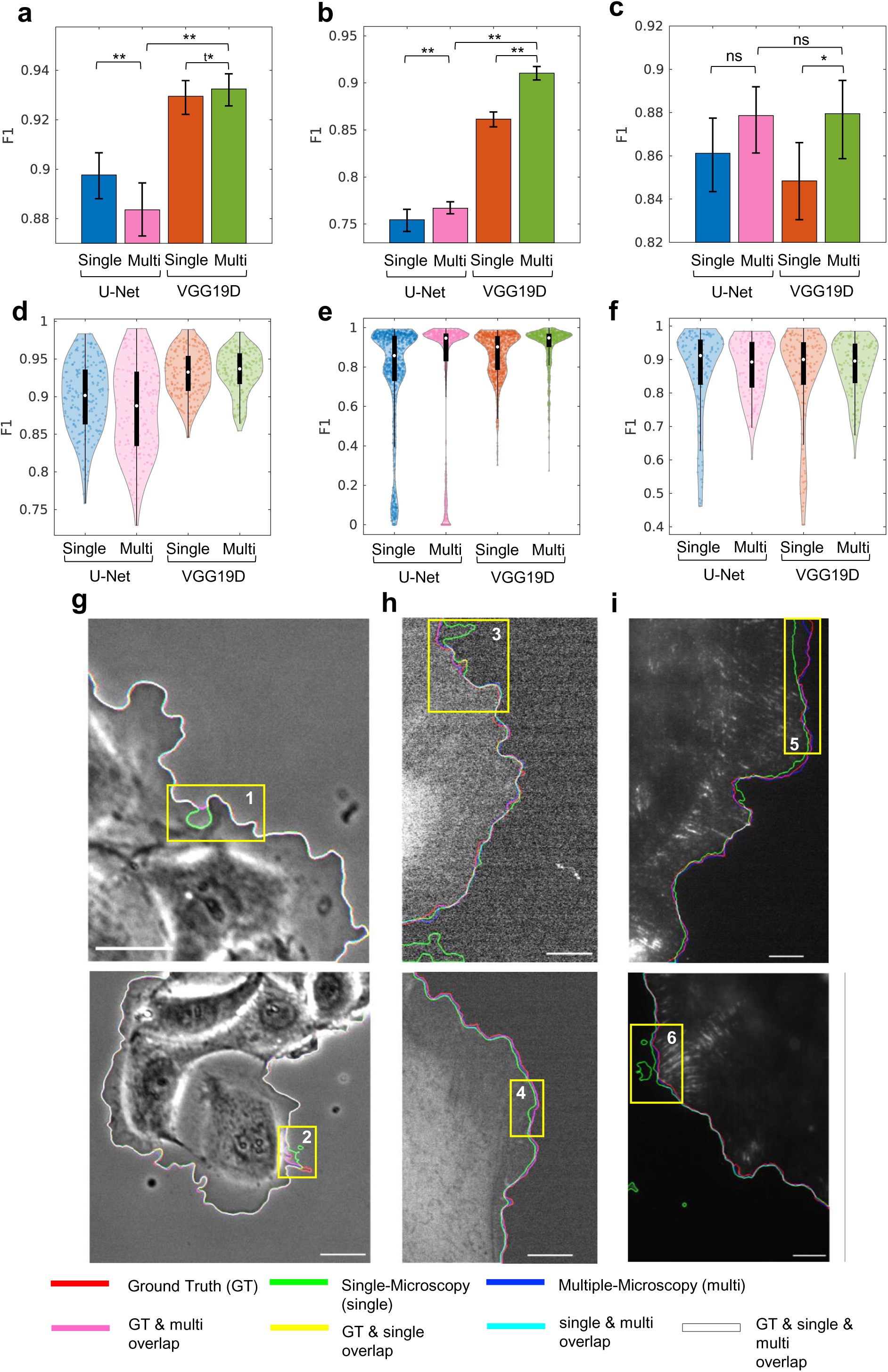
Comparisons of single-microscopy-type and multiple- microscopy-type training using U-Net and VGG19D-U-Net models. (**a-c**) Average F1 scores of models evaluated on phase contrast (**a**) , SDC (**b**) , and TIRF (**c**). (**d-f**) The distribution of F1 score in violin plot and box plot in black with a median indicated by the white circle for phase contrast (**d**) , SDC (**e**) , and TIRF (**f**) . The numbers of evaluated frames are n=202 for phase contrast, n=1000 for SDC, and n=132 for TIRF. The statistical significance by Wilcoxon signed-rank test with p >= 0.05 are indicated by ns, p <0.05 are indicated by *, and p < 0.001 are indicated by **. The statistical significance by paired t-test with p <0.05 is indicated by ^t^*. Error bars: 95% confidence intervals of the bootstrap mean. (**g-i**) Visualization of segmentation results on three microscopies: phase contrast (**g**), SDC (**h**) , and TIRF (**i**) . These correspond to the close-up views shown in Figure 6. Edges extracted from ground truth masks and predictions are overlaid on the original image. Each edge is represented by one of three primary colors. Overlap of two or more edges is represented by the combination of those colors. (**g**) Bar: 32.5 μm. (**h**) Bar: 7.2 μm. (**i**) Bar: 6.5 μm,

**Supplementary Figure 3.**
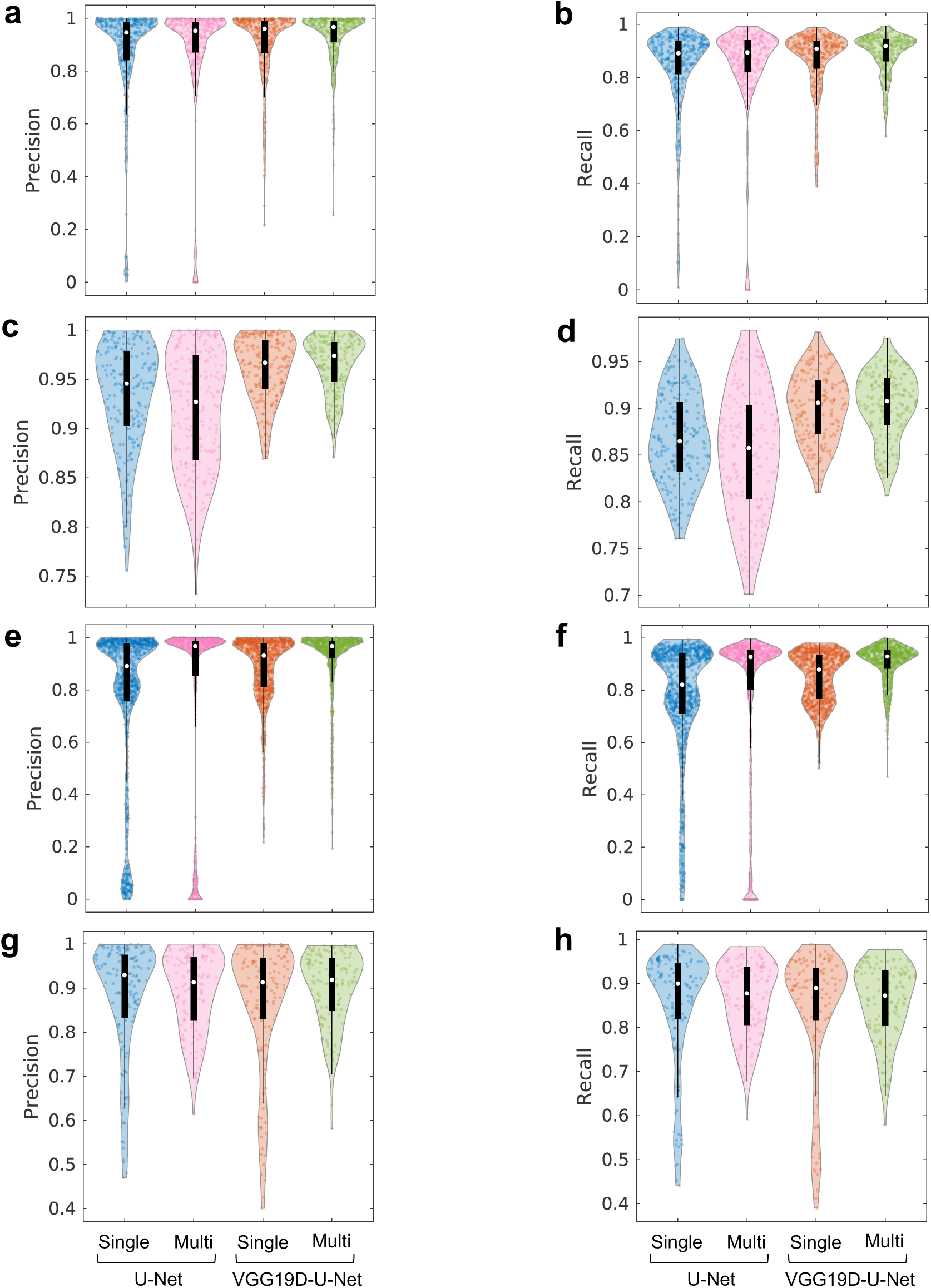
Comparisons of U-Net and VGG19D-U-Net models trained on single or multiple-microscopy-type dataset. (**a-h**) The distribution of precision and recall of U-Net and VGG19D-U-Net either trained on single-microscopy-type or multiple-microscopy- type datasets. The evaluated frames are sampled from all datasets (**a-b**) or phase contrast (**c-d)**, SDC (**e-f**) , or TIRF (**g-h**) dataset only. Within a violin plot, there is a box plot in black with a median indicated by the white circle. (n=335 for all datasets, n=202 for phase contrast, n=1000 for SDC, and n=132 for TIRF)

**Supplementary Figure 4.**
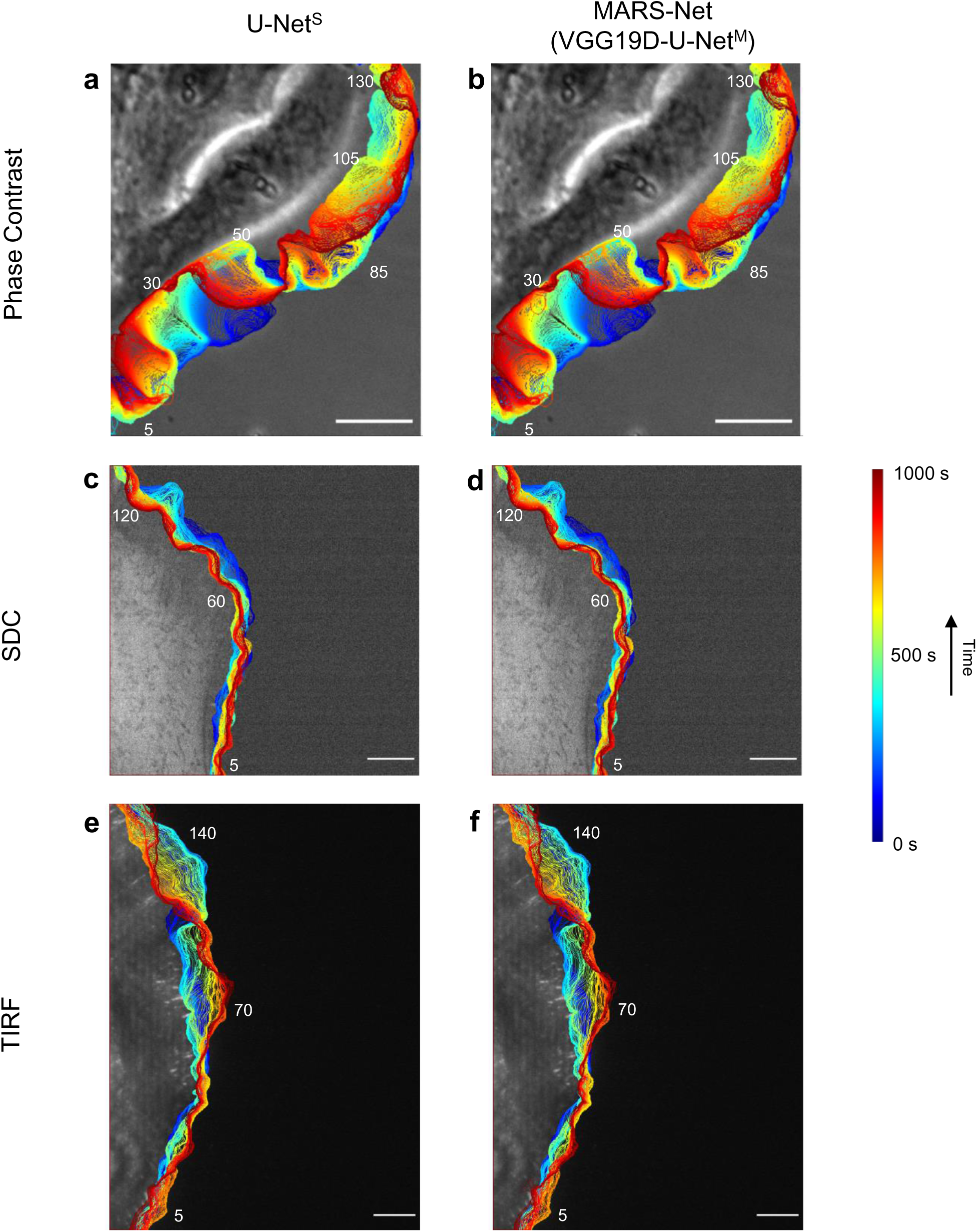
Visualization of single-microscopy-type U-Net and MARS-Net. (**a-f**) Progression of the cell edges overlaid on the first frame of the movie using segmentation results from U-Net^S^ (**a,c,e**) and MARS-Net (**b,d,f**). (**a-b**) Phase contrast live cell movie, Bars: 32.5 μm. (**c-d**) SDC live cell movie, Bars: 7.2 μm. (**e-f**) TIRF live cell movie, Bars: 6.5 μm. The approximate window numbers corresponding to window numbers in protrusion maps in Fig. 7 are labeled in white color. (blue, 0s; red, 1000s time points).

**Supplementary Figure 5.**
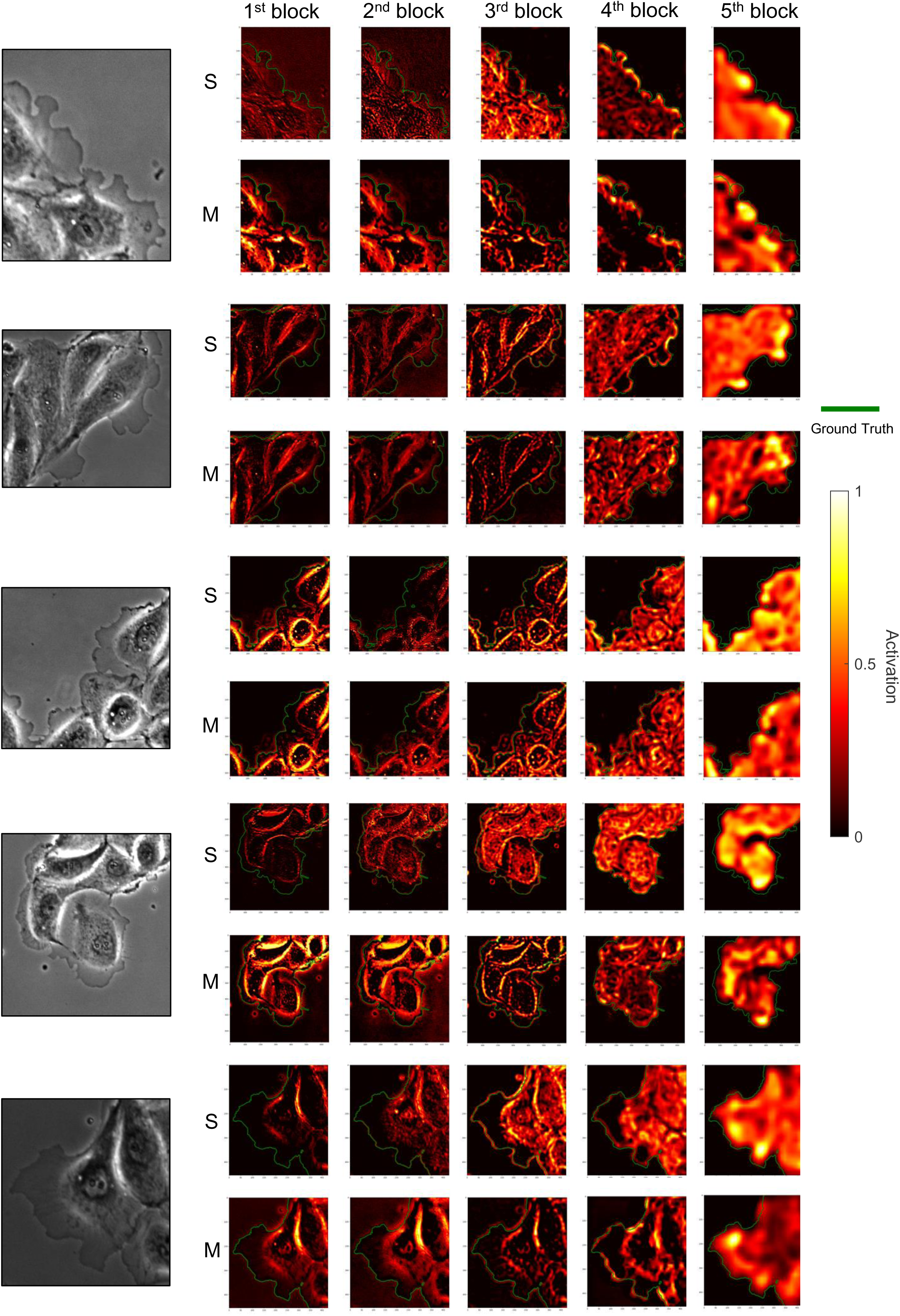
Class activation map of convolutional layers in the single-microscopy-type and multiple-microscopy-type VGG19D-U-Net with respect to the ground truth edges for the phase contrast dataset. From each live cell movie, one frame was randomly chosen for visualization. On the left, the original image of the corresponding frame is shown. The word ’S’ represents the single- microscopy-type dataset, and ’M’ represents the multiple-microscopy-type dataset. The class activation maps from the first to the fifth block in the encoder are organized in order from left to right. In the heatmap, the value 0 means no activation, and 1 means the highest activation. The green lines represent the ground truth edges.

**Supplementary Figure 6.**
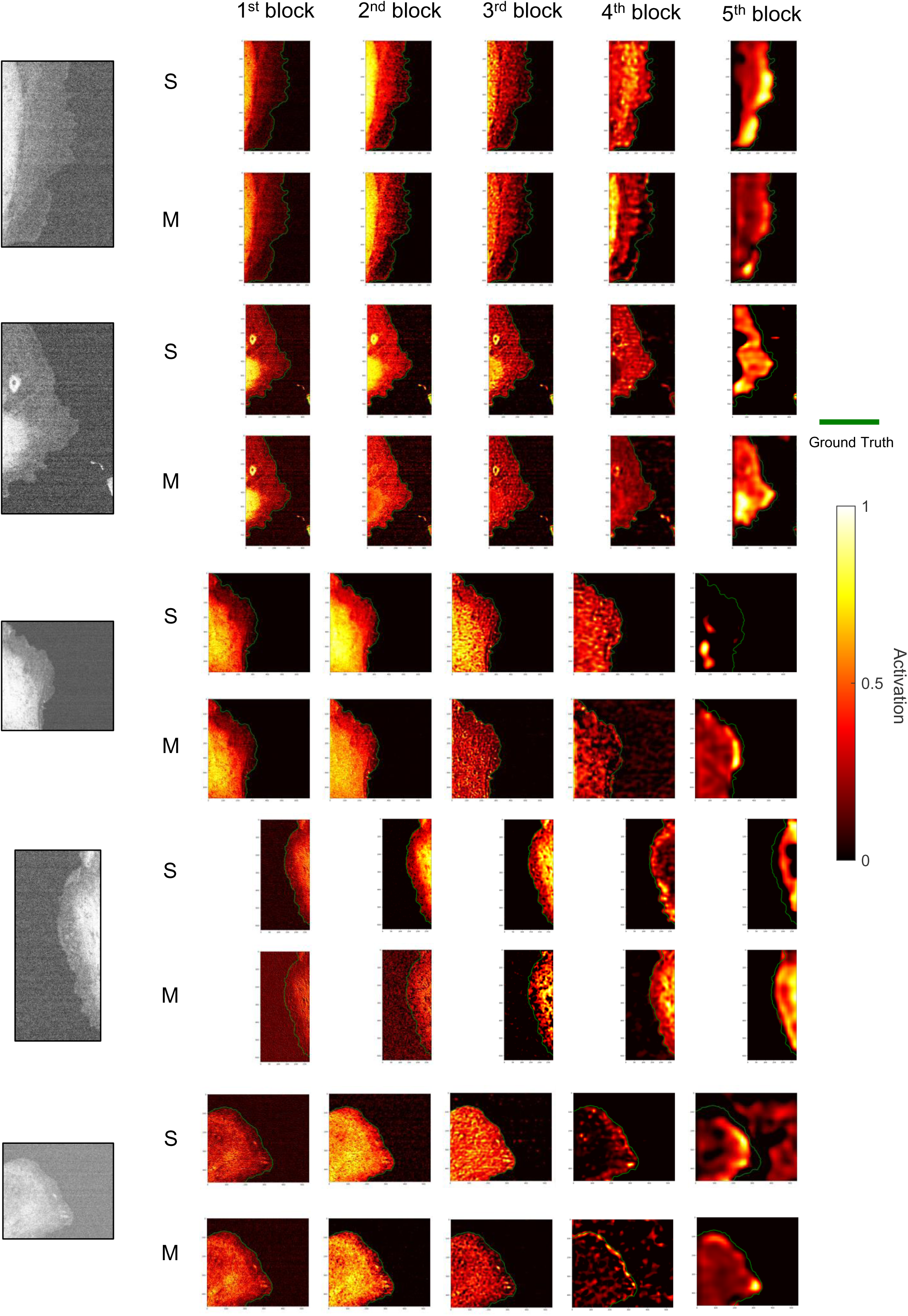
Class activation map of convolutional layers in the single-microscopy-type and multiple-microscopy-type VGG19D-U-Net with respect to the ground truth edges for the SDC dataset. From each live cell movie, one frame was randomly chosen for visualization. On the left, the original image of the corresponding frame is shown. The word ’S’ represents the single-microscopy-type dataset, and ’M’ represents the multiple-microscopy-type dataset. The class activation maps from the first to the fifth block in the encoder are organized in order from left to right. In the heatmap, the value 0 means no activation, and 1 means the highest activation. The green lines represent the ground truth edges.

**Supplementary Figure 7.**
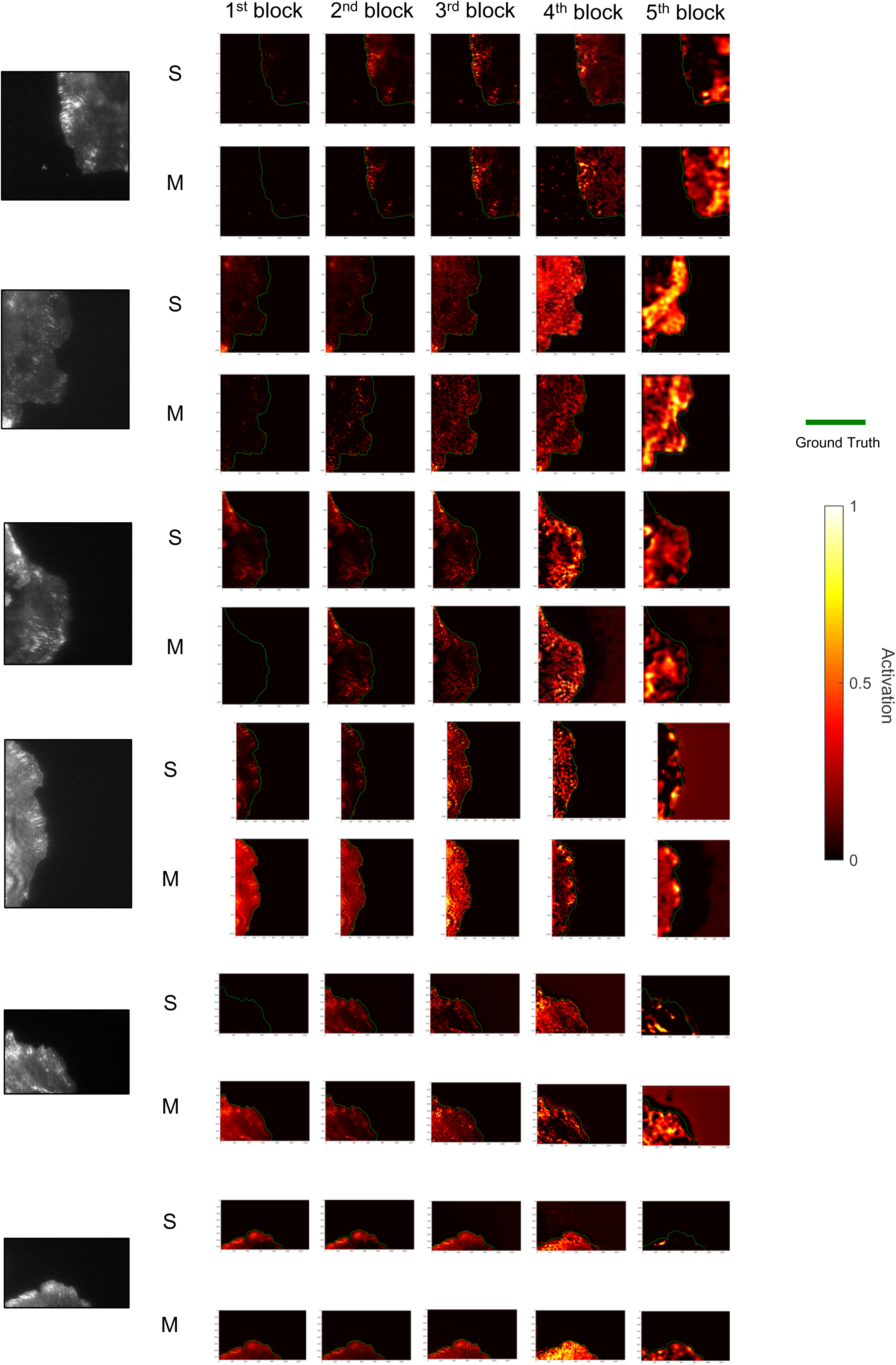
Class activation map of convolutional layers in the single-microscopy-type and multiple-microscopy-type VGG19D-U-Net with respect to the ground truth edge for the TIRF dataset. From each live cell movie, one frame was randomly chosen for visualization. On the left, the original image of the corresponding frame is shown. The word ’S’ represents the single-microscopy-type dataset, and ’M’ represents the multiple-microscopy-type dataset. The class activation maps from the first to the fifth block in the encoder are organized in order from left to right. In the heatmap, the value 0 means no activation, and 1 means the highest activation. The green lines represent the ground truth edges.

**Supplementary Figure 8.**
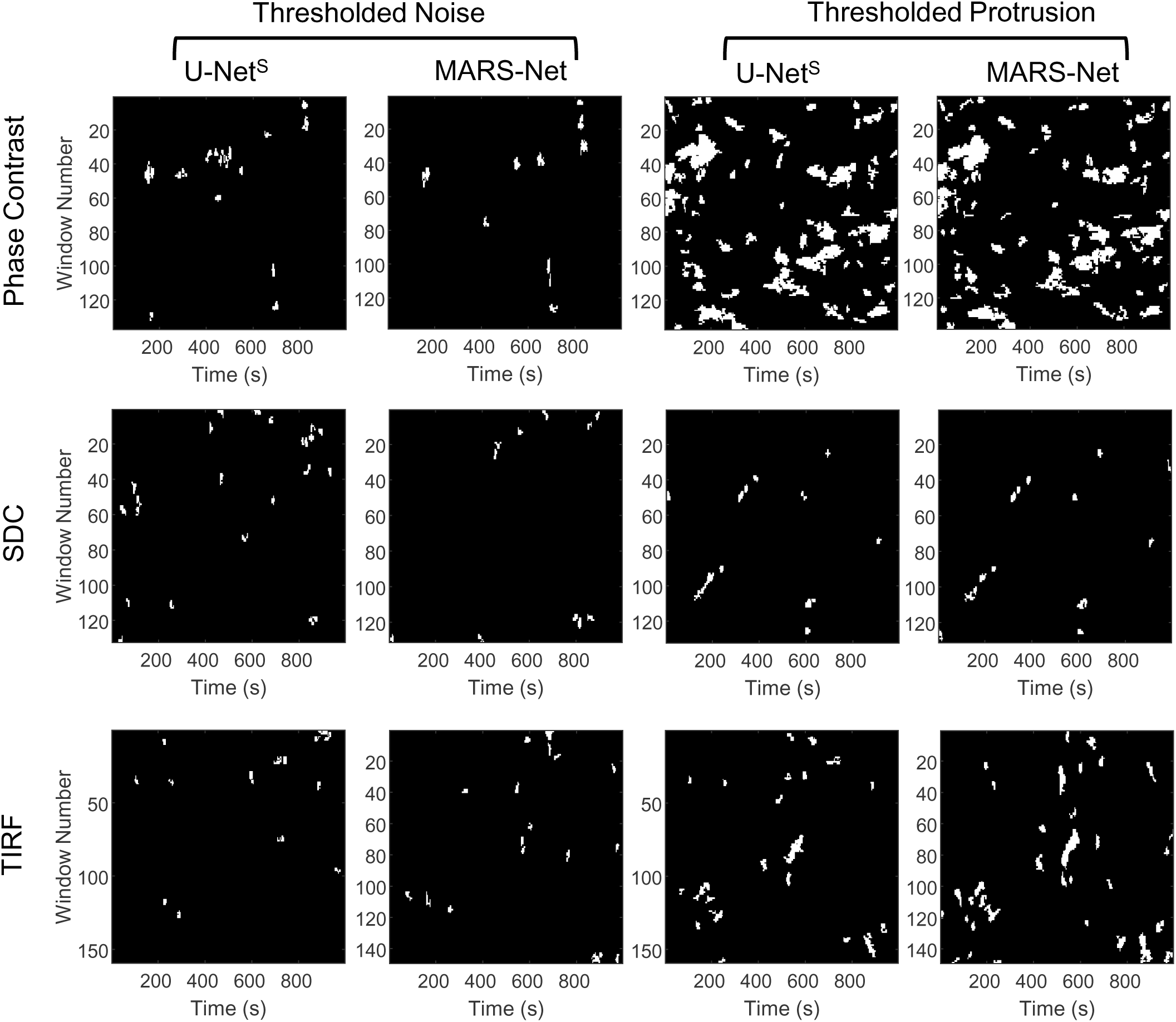
Threshold error/noise and protrusion in the protrusion maps of Phase Contrast, SDC, and TIRF live cell movies segmented by single-microscopy-type U-Net and MARS-Net. These are obtained from filtering and thresholding protrusion maps in Fig. 7. In each row, the same cropped region in the movie is profiled for cellular morphodynamics.

